# Functional Annotation of Human Long Non-Coding RNAs via Molecular Phenotyping

**DOI:** 10.1101/700864

**Authors:** Jordan A Ramilowski, Chi Wai Yip, Saumya Agrawal, Jen-Chien Chang, Yari Ciani, Ivan V Kulakovskiy, Mickaël Mendez, Jasmine Li Ching Ooi, John F Ouyang, Nick Parkinson, Andreas Petri, Leonie Roos, Jessica Severin, Kayoko Yasuzawa, Imad Abugessaisa, Altuna Akalin, Ivan V Antonov, Erik Arner, Alessandro Bonetti, Hidemasa Bono, Beatrice Borsari, Frank Brombacher, Chris JF Cameron, Carlo Vittorio Cannistraci, Ryan Cardenas, Melissa Cardon, Howard Chang, Josée Dostie, Luca Ducoli, Alexander Favorov, Alexandre Fort, Diego Garrido, Noa Gil, Juliette Gimenez, Reto Guler, Lusy Handoko, Jayson Harshbarger, Akira Hasegawa, Yuki Hasegawa, Kosuke Hashimoto, Norihito Hayatsu, Peter Heutink, Tetsuro Hirose, Eddie L Imada, Masayoshi Itoh, Bogumil Kaczkowski, Aditi Kanhere, Emily Kawabata, Hideya Kawaji, Tsugumi Kawashima, S. Thomas Kelly, Miki Kojima, Naoto Kondo, Haruhiko Koseki, Tsukasa Kouno, Anton Kratz, Mariola Kurowska-Stolarska, Andrew Tae Jun Kwon, Jeffrey Leek, Andreas Lennartsson, Marina Lizio, Fernando López-Redondo, Joachim Luginbühl, Shiori Maeda, Vsevolod J Makeev, Luigi Marchionni, Yulia A Medvedeva, Aki Minoda, Ferenc Müller, Manuel Muñoz-Aguirre, Mitsuyoshi Murata, Hiromi Nishiyori, Kazuhiro R Nitta, Shuhei Noguchi, Yukihiko Noro, Ramil Nurtdinov, Yasushi Okazaki, Valerio Orlando, Denis Paquette, Callum JC Parr, Owen JL Rackham, Patrizia Rizzu, Diego Fernando Sánchez Martinez, Albin Sandelin, Pillay Sanjana, Colin AM Semple, Youtaro Shibayama, Divya M Sivaraman, Takahiro Suzuki, Suzannah C Szumowski, Michihira Tagami, Martin S Taylor, Chikashi Terao, Malte Thodberg, Supat Thongjuea, Vidisha Tripathi, Igor Ulitsky, Roberto Verardo, Ilya Vorontsov, Chinatsu Yamamoto, Robert S Young, J Kenneth Baillie, Alistair RR Forrest, Roderic Guigó, Michael M Hoffman, Chung Chau Hon, Takeya Kasukawa, Sakari Kauppinen, Juha Kere, Boris Lenhard, Claudio Schneider, Harukazu Suzuki, Ken Yagi, FANTOM consortium, Michiel de Hoon, Jay W Shin, Piero Carninci

## Abstract

Long non-coding RNAs (lncRNAs) constitute the majority of transcripts in the mammalian genomes and yet, their functions remain largely unknown. We systematically knockdown 285 lncRNAs expression in human dermal fibroblasts and quantified cellular growth, morphological changes, and transcriptomic responses using Capped Analysis of Gene Expression (CAGE). Antisense oligonucleotides targeting the same lncRNA exhibited global concordance, and the molecular phenotype, measured by CAGE, recapitulated the observed cellular phenotypes while providing additional insights on the affected genes and pathways. Here, we disseminate the largest to-date lncRNA knockdown dataset with molecular phenotyping (over 1,000 CAGE deep-sequencing libraries) for further exploration and highlight functional roles for *ZNF213-AS1* and *lnc-KHDC3L-2.*

## Introduction

Over 50,000 loci in the human genome transcribe long non-coding RNA (lncRNA) (Hon et al. 2017; Iyer et al. 2015), which are defined as transcripts at least 200 nucleotides long with low or no protein-coding potential. While lncRNA genes outnumber protein-coding genes in mammalian genomes, they are comparatively less conserved (Ulitsky 2016), lowly expressed, and more cell-type-specific (Hon et al. 2017). However, the evolutionary conservation of lncRNA promoters (Carninci et al. 2005) and their structural motifs of lncRNAs (Xue et al. 2016), (Chu et al. 2015) suggest that lncRNAs are fundamental biological regulators. To date, only a few hundred human lncRNAs have been extensively characterized (Quek et al. 2015; Volders et al. 2015; de Hoon et al. 2015; Ma et al. 2019), revealing their roles in regulating transcription (Engreitz, Ollikainen, et al. 2016), translation (Carrieri et al. 2012), and chromatin state (Gupta et al. 2010; Guttman and Rinn 2012; Guttman et al. 2011); (Ransohoff et al. 2018; Quinn and Chang 2016).

Our recent FANTOM 5 computational analysis showed that 19,175 (out of 27,919) human lncRNA loci are functionally implicated (Hon et al. 2017). Yet, genomic screens are necessary to comprehensively characterize each lncRNA. One common approach of gene knockdown followed by a cellular phenotype assay typically characterizes a small percentage of lncRNAs for a single observable phenotype. For example, a recent large-scale screening using CRISPR interference (CRISPRi) found that approximately ~3.7% of targeted lncRNA loci are essential for cell growth or viability in a cell-type specific manner (Liu et al. 2017). In addition, CRISPR-Cas9 experiments targeting splice sites identified ~2.1% of lncRNAs that affect growth of K562 (Liu et al. 2018) and a CRISPR activation study revealed ~0.11% lncRNAs to be important for drug resistance in melanoma (Joung et al. 2017). However, many of these studies target the genomic DNA, potentially perturbing the chromatin architecture, or focus on a single cellular assay, possibly missing other relevant functions and underlying molecular pathways.

As a part of the FANTOM 6 pilot project, we established an automated high-throughput cell culture platform to suppress 285 lncRNAs expressed in human primary dermal fibroblasts (HDF) using antisense LNA-modified GapmeR antisense oligonucleotide (ASO) technology (Roux et al. 2017). We then quantified the effect of each knockdown on cell growth and morphology using real-time imaging, followed by Cap Analysis Gene Expression (CAGE; (Murata et al. 2014) deep sequencing to reveal molecular pathways associated with each lncRNA. In contrast to cellular phenotyping, molecular phenotyping provides a detailed assessment of the response to an lncRNA knockdown at the molecular level, allowing biological pathways to be associated to lncRNAs even in the absence of an observable cellular phenotype. All data and analysis results are publicly available (see Data Access) and results can be interactively explored using our inhouse portal https://fantom.gsc.riken.jp/zenbu/reports/#FANTOM6/.

## Results

### Selection and ASO-mediated knockdown of lncRNA targets

Human dermal fibroblasts (HDF) are non-transformed primary cells that are commonly used for investigating cellular reprogramming (Takahashi et al. 2007; Ambasudhan et al. 2011), woundhealing (Li and Wang 2011), fibrosis (Kendall R., et al 2014), and cancer (Kalluri 2016). Here, an unbiased selection of lncRNAs expressed in HDF was performed to choose 285 lncRNAs for functional interrogation (Methods; Supplemental Table S1, Fig. 1A-C). Using RNA-seq profiling of fractionated RNA, we annotated the lncRNA subcellular localization in the chromatin-bound (35%), nucleus-soluble (27%), or cytoplasm (38%) (Fig. 1D). We then designed a minimum of five non-overlapping antisense oligonucleotides (ASOs) against each lncRNA (Supplemental Methods; Supplemental Table S2; Fig. 1E,F) and transfected them individually using an automated cell culture platform to minimize experimental variability (Fig. 1G). The overall knockdown efficiencies across 2,021 ASOs resulted in median value of 45.4%, and we could successfully knockdown 879 out of 2,021 (43.5%) ASOs (>40% knockdown efficiency in at least two primer pairs or >60% in one primer pair; Supplemental Table S2). ASOs targeting exons or introns were equally effective, and knockdown efficiencies were independent of the genomic class, expression level, and subcellular localization of the lncRNA (Supplemental Fig. S1A–D).

**Figure 1.**
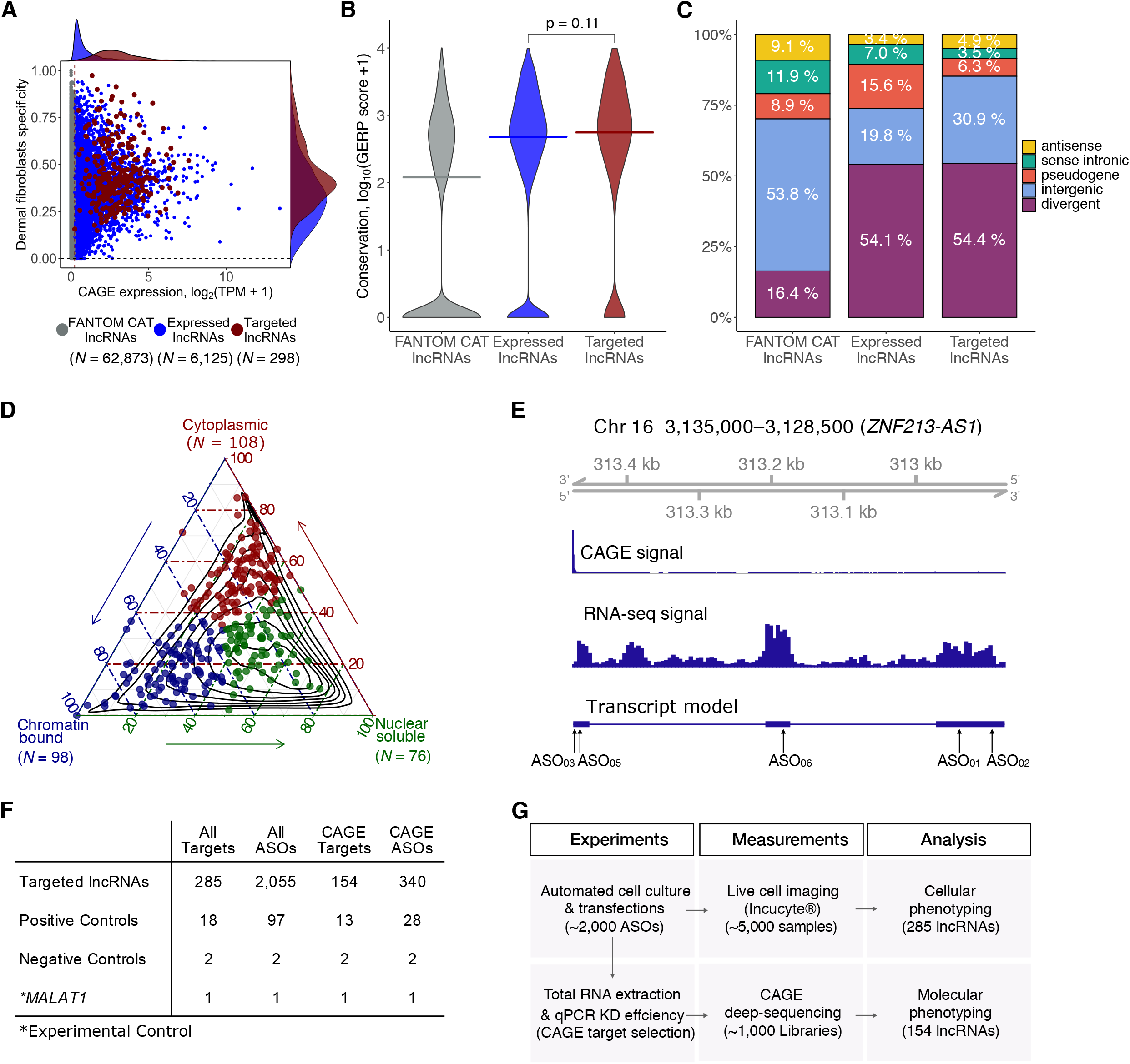
Selection of lncRNA targets, their properties and the study overview. (*A*) CAGE expression levels at log2TPM (tags per million) and human dermal fibroblasts (HDF) specificity of lncRNAs in the FANTOM CAT catalog (Hon, et al., Nature 2017; *N* = 62,873; grey), lncRNAs expressed in HDF (*N* = 6,125; blue) and targeted lncRNAs (*N* = 285; red). The dashed vertical line indicates most lowly expressed lncRNA target (~0.2 TPM). (*B*) Gene conservation levels of lncRNAs in the FANTOM CAT catalog (grey), lncRNAs expressed in HDF (blue) and targeted lncRNAs (red). Crossbars indicate the median. No significant difference is observed when comparing targeted and expressed in HDF lncRNAs (Wilcoxon p = 0.11). (*C*) Similar to that in B, but for genomic classes of lncRNAs. Most of the targeted lncRNAs and those expressed in HDF are expressed from divergent promoters. (*D*) Subcellular localization (based on relative abundances from RNA-seq fractionation data) for targeted lncRNAs. Chromatin-bound (*N* = 98; blue); Nuclear soluble (*N* = 76; green); Cytoplasmic (*N* = 108; red). Black contours represent the distribution of all lncRNAs expressed in HDF. (*E*) Example of *ZNF213-AS1* loci showing transcript model, CAGE and RNA-seq signal along with targeting ASOs. (*F*) Number of ASOs for target lncRNAs and controls used in the experiment. (*G*) Schematics of the study.

### A subset of lncRNAs associated with cell growth and morphology changes

To evaluate the effect of each lncRNA knockdown on cell growth and morphology, we imaged ASO-transfected HDF in duplicates every 3 hours for a total of 48 hours (Supplemental Table S3) and estimated their growth rate based on cell confluence measurements (Fig. 2A,B). First, we observed across all ASOs that changes in cell growth and morphological parameters were significantly correlated with knockdown efficiency (Supplemental Fig. S1E). Considering both successful knockdown and significant growth inhibition (Student’s two-sided *t*-test FDR ≤ 0.05), 246 out of 879 ASOs (~28%) showed cellular phenotype (Fig. 2C, Table S3).

**Figure 2.**
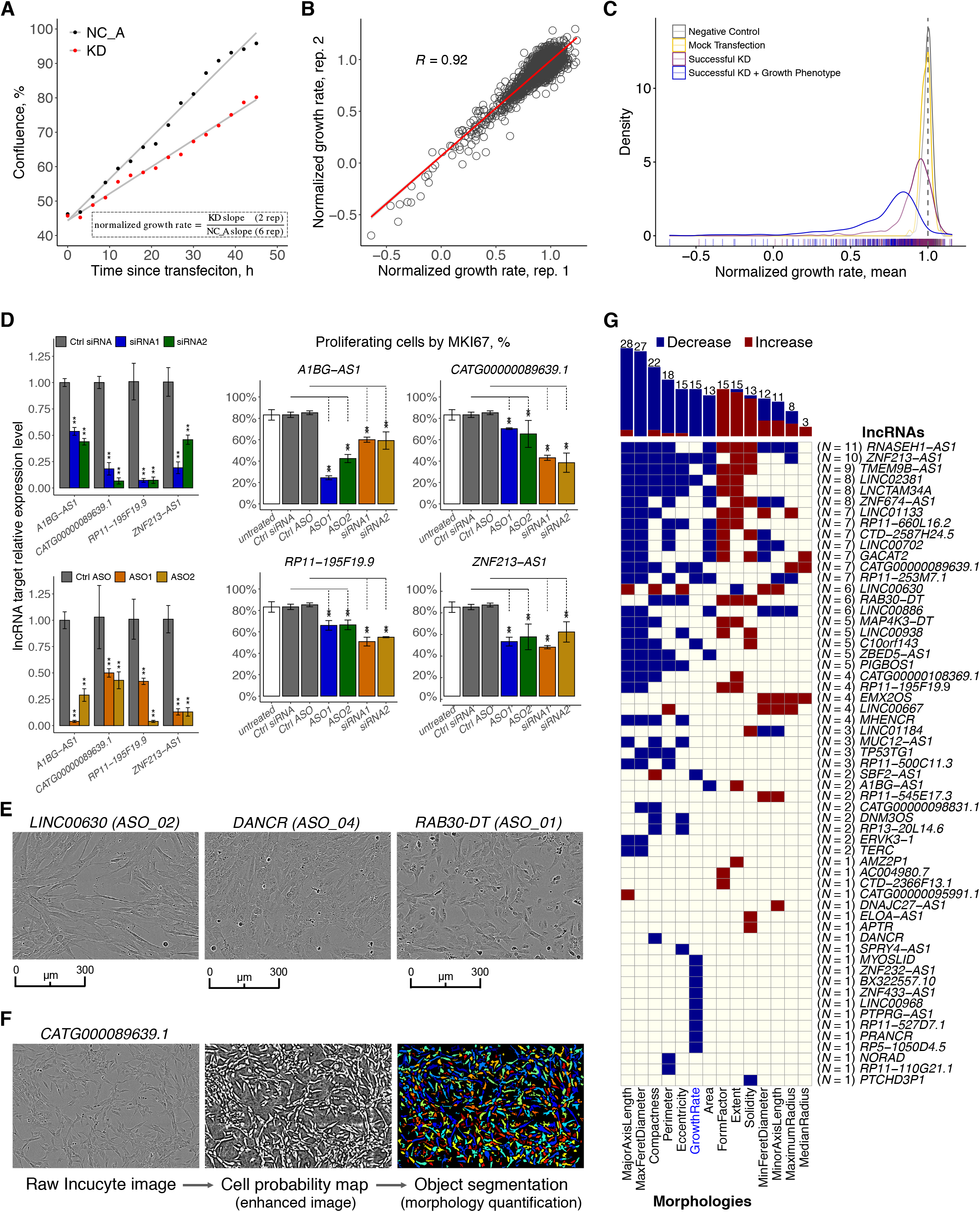
Cell growth and morphology assessment. (*A*) Selected example *(PTPRG1-AS1)* showing the normalized growth rate estimation using a matching NC_A (negative control). (*B*) Correlation of the normalized growth rate for technical duplicates across 2,456 IncuCyte^®^ samples. (*C*) Density distribution of normalized growth rates (technical replicates averaged) 252 ASOs targeting lncRNAs with successful knockdown (KD) and growth phenotype (blue) consistent in 2 replicates (FDR < 0.05 as compared to matching NC_A; 246 ASOs inhibited growth), 627 ASOs targeting lncRNAs with successful KD (purple), 270 negative control (NC_A) samples (grey) and 90 mock-transfected cells (lipofectamine only) samples (yellow). (*D*) MKI67 staining (growth inhibition validation) for four selected lncRNA targets after siRNA and ASOs suppression. *(E)* IncuCyte^®^ cell images of selected distinct cell morphologies changes upon an lncRNA KD. (*F*) An overview of cell morphology imaging processing pipeline using a novel lncRNA target, *CATG000089639.1,* as an example. (*G*) lncRNAs (*N* = 59) significantly (FDR < 0.05) and consistently (after adjusting for the number of successfully targeting ASOs) affecting cell growth (*N* = 15) and cell morphologies (*N* = 44).

To assess globally whether the observed growth inhibition is lncRNA-specific, we used all 194 lncRNAs successfully targeted by at least two ASOs (Supplemental Fig. S2A) and found that ASOs targeting the same lncRNA were significantly more likely to have a concordant growth response than ASOs targeting different lncRNA (empirical p = 0.00037; Supplemental Methods; Supplemental Fig. S2B). However, different ASOs targeting the same lncRNA typically showed different effects on growth, possibly due to variable knockdown efficiencies, differences in targeted lncRNA isoforms, as well as off-target effects. To reliably identify target specific cellular phenotype, we applied conditional cutoffs based on the number of successful ASOs per each lncRNA (Supplemental Methods; Supplemental Fig. S2C) and identified 15/194 lncRNAs (7.7%) with growth phenotype (adjusted background less than 5%; Supplemental Fig. S2D). We validated *A1BG-AS1,* which was previously implicated in cell growth (Bai et al. 2019), *CATG00000089639, RP11-195F19.9,* and *ZNF213-AS1* by measuring the MKI67 proliferation protein marker upon knockdown with siRNAs and selected ASOs (Fig. 2D, Supplemental Fig. S2E).

In addition to cell growth, we also explored changes in cell morphology (Fig. 2E). Using a machine learning-assisted workflow (Methods), each cell was segmented and its morphological features representing various aspects of cell shapes and sizes were quantified (Carpenter et al. 2006) (Fig. 2F; Supplemental Table S3). As an example, knockdown of 14/194 lncRNAs (7.2%) affected the spindle-like morphology of fibroblasts, as indicated by a consistent decrease in their observed eccentricity without reducing the cell number, suggesting possible cellular transformation towards epithelial-like states. Collectively, we observed 59/194 lncRNAs (~30%) affecting cell growth and/or morphological parameters (Fig. 2G; Supplemental Table S3)

### Molecular phenotyping by CAGE recapitulates cellular phenotypes and highlights functions of lncRNAs

Next, we selected 340 ASOs with high knockdown efficiencies (mostly greater than 50%; median 71.4%) and sequenced 970 CAGE libraries to analyze 154 lncRNAs (Fig. 3A; Supplemental Table S4). To assess functional implications by individual ASOs, we performed differential gene expression, Motif Activity Response Analysis (MARA; (FANTOM Consortium et al. 2009), and Gene Set Enrichment Analysis (GSEA; (Subramanian et al. 2005); Fig 3B-E), and compared them with cellular phenotype.

**Figure 3.**
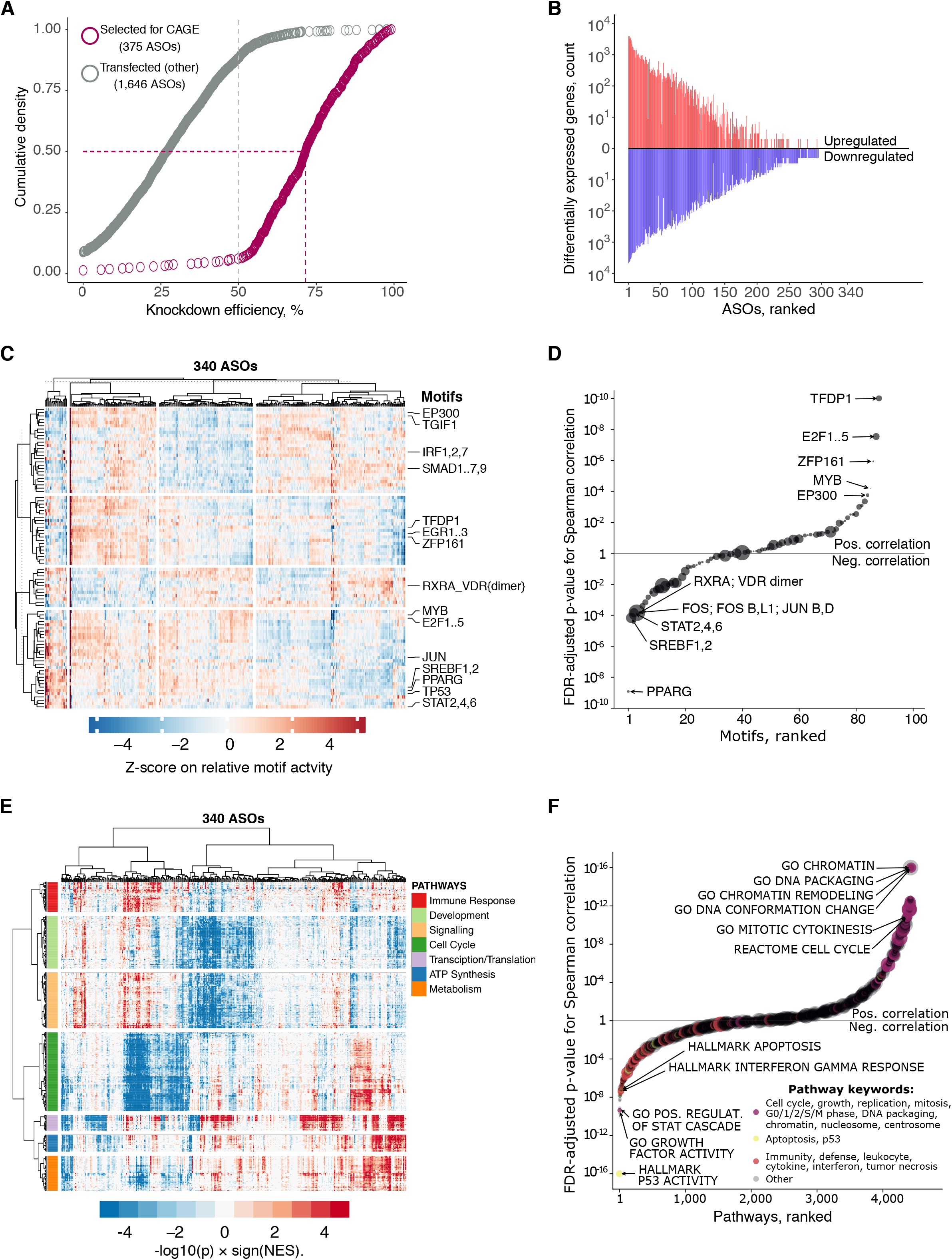
CAGE predicts cellular phenotypes. (*A*) RT-qPCR knockdown (KD) efficiency for 2,021 ASO-transfected samples (targeted lncRNAs only). Grey dashed line indicates 50% KD efficiency generally required for CAGE selection. Purple dashed lines indicate median KD efficiency (71.5%) for 375 ASOs selected for CAGE sequencing. After quality control, 340 ASOs targeting lncRNAs were included for further analysis. (*B*) Distribution of significantly differentially expressed genes (up-regulated: FDR < 0.05, Z-score > 1.645, log2FC > 0.5 and down-regulated: FDR < 0.05, Z-score < −1.645, log2FC < −0.5) across all 340 ASOs. (*C*) Motif Response Activity Analysis (MARA) across 340 ASOs. Scale indicates Z-score of the relative motif activity (the range was set to abs(Z-score) = < 5 for visualization purposes). *(D)* Correlation between normalized growth rate and motif activities across 340 ASOs targeting lncRNAs with highlighted examples. Motifs sizes shown are scaled based on the HDF expression of their associated TFs (range 1 to ~600 TPM). (*E*) Enriched biological pathways across 340 ASOs. Scale indicates GSEA enrichment value calculated as -log10(p) × sign(NES). (*F*) same as in D, but for selected GSEA pathways. Pathways sizes are scaled based on the number of associated genes.

We globally observed significant knockdown-mediated transcriptomic changes (which generally correlated with KD efficiency; Supplemental Fig. S3A), with ~57% of ASOs showing at least 10 differentially expressed genes (FDR ≤ 0.05; abs(log_2_FC) > 0.5). For 84 divergent-antisense lncRNAs (targeted by 186 independent ASOs) (Supplemental Methods), we found their partner gene to be generally unchanged (median abs(log_2_FC) = ~0.13), with an exception of two significantly downregulated and three significantly upregulated genes (FDR ≤ 0.05; Supplemental Fig. S3B). We have, however, noticed common response in a large number of ASOs (~30-35% of all responding ASOs) such as down-regulation of cell-cycle related pathways, upregulated stress genes and pathways or altered cell metabolism and energetics (Supplemental Fig. S3C,D).

When comparing knockdown-mediated molecular and cellular response, we found that transcription factor motifs that promote cell growth, including TFDP1, E2F1,2,3, and EP300, were positively correlated with the measured cell growth rate while transcription factor motifs known to inhibit growth or induce apoptosis *(e.g.* PPARG, SREBPF, and STAT2,4,6) were negatively correlated (Fig. 3D; Supplemental Fig. S4A; Supplemental Table S6). Moreover, correlations between GSEA pathways (Fig. 3F; Supplemental Fig. S4B; Supplemental Table S6) and FANTOM5 co-expression clusters (Supplemental Fig. S4C) showed that cell growth and replication related pathways were positively correlated with the measured growth rate, whereas those related to immunity, cell stress and cell death were negatively correlated. We found that amongst 53 ASOs implicated in growth inhibition pathway based on the CAGE profiles, only 43% of them showed growth inhibition in the real-time imaging. This might suggest better sensitivity of transcriptomic profiling when detecting phenotypes as compared to live cell imaging methods, which are more prone to a delayed cellular response to the knockdown.

Additionally, morphological changes were reflected in the molecular phenotype assessed by CAGE (Supplemental Fig. S4D). Cell radius and axis length were associated with GSEA categories related to actin arrangement and cilia, while cell compactness was negatively correlated with apoptosis. The extensive molecular phenotyping analysis also revealed pathways not explicitly associated with cell growth and cell morphology, such as transcription, translation, metabolism, development and signaling (Fig. 3E).

Next, to globally assess whether individual ASO knockdowns lead to lncRNA-specific effects, we scaled the expression change of each gene across the whole experiment and compared differentially expressed genes (Fig. 3B) of all possible ASO pairs targeting the same lncRNA target versus different lncRNAs (Supplemental Methods; Supplemental Table S5). We found that the concordance of the same target group was significantly greater than that of the different target group (comparing the Jaccard indices across 10,000 permutations; Supplemental Fig. S5A), suggesting that ASO knockdowns are non-random and lead to more lncRNA specific effects than the non-targeting ASO pairs. Further, by requiring at least five common DEGs (FDR ≤ 0.05, abs(log_2_FC) > 0.5, abs(Z-score) > 1.645) and ASO-pairs significantly above the nontargeting ASO pairs background (p ≤ 0.05), we identified 16 ASO-pairs, targeting 13 lncRNAs, exhibiting reproducible knockdown-mediated molecular responses in human dermal fibroblasts (Supplemental Fig. S5B). Corresponding GSEA pathways and MARA motifs of these 16 ASO-pairs are shown in Supplemental Figure S5C.

### siRNA validation experiments

To evaluate whether the lncRNA-specific effects can be measured by other knockdown technologies, nine lncRNAs, with relatively mild growth phenotype, were subjected to siRNA knockdown. We noted that higher concordance was observed for ASO modality alone (Supplemental Fig. S5D). The observed discrepancies in the transcriptional response between ASO and siRNA-mediated knockdowns could be contributed by their mode of action and variable activities in different subcellular compartments. Next, a concordant response was found for (5/36) ASO-siRNA pairs targeting three lncRNAs (Supplemental Fig. S5E; Supplemental Table S5), enriched in the cytoplasm *(MAPKAPK5-AS1),* soluble nuclear fraction *(LINC02454)* and in the chromatin-bound fraction (*A1BG-AS1*). While we cannot completely exclude the technical artefacts of each technology, concordant cellular response exhibited by using ASOs alone suggests that lncRNA, in part, are essential regulatory elements in cells. Yet, our study generally warrants a careful assessment of specific findings from different knockdown technologies, including CRISPR-inhibition, and demonstrates a requirement of using multiple replicates in a given target per each modality.

### *ZNF213-AS1* is associated with cell growth and migration

Extensive molecular and cellular phenotype data for each ASO knockdown can be explored using our portal https://fantom.gsc.riken.jp/zenbu/reports/#FANTOM6. As an example of an lncRNA associated with cell growth and morphology (Fig. 2G), we showcase *ZNF213-AS1 (RP11-473M20.14*\ This lncRNA is highly conserved in placental mammals, moderately expressed (~8 CAGE tags per million) in HDF and enriched in the chromatin-bound fraction. Four distinct ASOs (ASO_01, ASO_02, ASO_05, and ASO_06) strongly suppressed expression of *ZNF213-AS1*, while expression of the *ZNF213* sense gene was not significantly affected in any of the knockdowns. The four ASOs caused varying degrees of cell growth inhibition (Fig. 4A). ASO_01 and ASO_06 showed a reduction in cell number, as well as an upregulation of apoptosis, immune and defense pathways in GSEA suggesting cell death. While cell growth inhibition observed for ASO_02 and ASO_05 was confirmed by MKI67 marker staining (Fig. 2D; Supplemental Tables S7), the molecular phenotype revealed suppression of GSEA pathways related to cell growth, as well as to cell proliferation, motility, and extracellular structure organization (Fig. 4B), and consistent in two ASOs downregulation of related motifs, for example, EGR1, EP300, SMAD1..7,9 (Fig. 4C).

**Figure 4.**
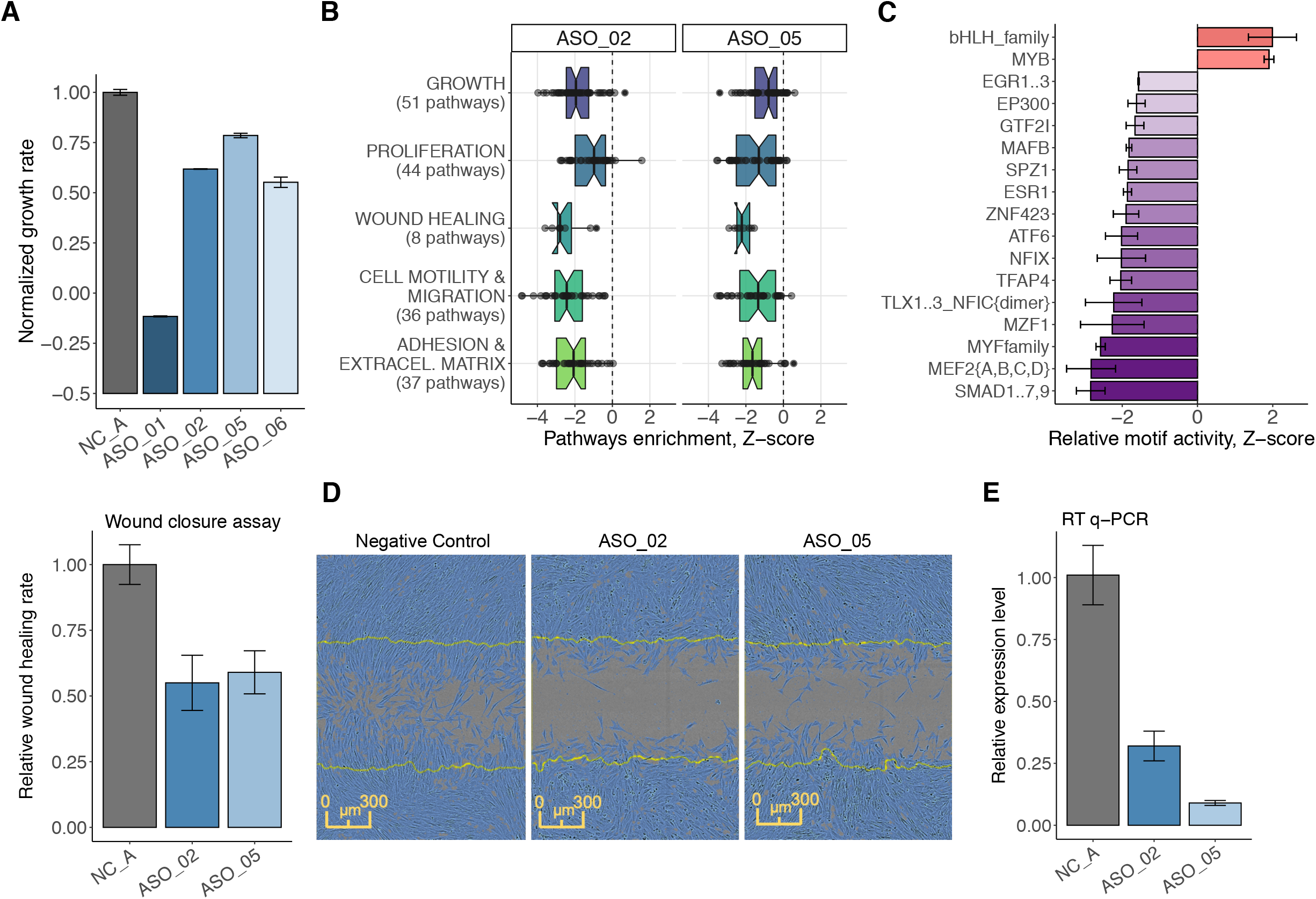
*ZNF213-AS1* regulates cell growth, migration and proliferation. (*A*) Normalized growth rate across four distinct ASOs (in duplicates) targeting *ZNF213-AS1* as compared to six negative control samples (shown in grey). (*B*) Enrichment of biological pathways associated with growth, proliferation, wound healing, migration and adhesion for ASO_02 and ASO_05. (*C*) Most consistently down- and upregulated transcription factor binding motifs including those for transcription factors known to modulate growth, migration and proliferation such as for example EGR family, EP300, GTF2I. (*D*) Transfected, re-plated and mitomycin-C (5 μg/mL)-treated HDF cells were scratched and monitored in the IncuCyte^®^ imaging system. Relative wound closure rate calculated during the 24 hours post-scratching shows 40-45% reduction for the two targeting ASOs (ASO_02 (*N* = 10) and ASO_05 (*N* = 13)) as compared to NC_A transfection controls (*N* = 33, shown in grey) and the representative images of wound closure assay 16 hours post-scratching. (*E*) Knockdown efficiency measured by RT-qPCR after wound closure assay (72 hours post-transfection) showing sustained suppression (65-90%) of *ZNF213-AS1*.

As cell motility pathways were affected by the knockdown, we tested whether *ZNF213-AS1* could influence cell migration. Based on the wound-closure assay after transient cell growth inhibition (mitomycin-C and serum starvation), we observed a substantial reduction of wound closure rate (~40% over a 24-hour period) in the *ZNF213-AS1* depleted HDFs (Fig. 4D,E). The reduced wound healing rate should thus mainly reflect reduced cell motility, further confirming affected motility pathways predicted by the molecular phenotype.

As these results indicated a potential role of *ZNF213-AS1* in cell growth and migration, we used FANTOM CAT Recount 2 atlas (Imada et al. 2020), which incorporates the TCGA dataset (Collado-Torres et al. 2017), and found relatively higher expression of *ZNF213-AS1* in acute myeloid leukemia (LAML) and in low-grade gliomas (LGG) as compared to other cancers (Supplemental Fig. S6A). In LAML, the highest expression levels were associated with mostly undifferentiated states, whereas in LGG, elevated expression levels were found in oligodendrogliomas, astrocytomas, and in IDH1 mutated tumors, suggesting that *ZNF213-AS1* is involved in modulating differentiation and proliferation of tumors (Supplemental Fig. S6B-E). Further, univariate Cox proportional hazard analysis as well as Kaplan-Meier curves for LGG were significant and consistent with our findings (HR = 0.61, BH FDR = 0.0079). The same survival analysis on LAML showed a weak association with poor prognostic outcome but the results were not significant; (Supplemental Fig. S6F,G).

### *RP11-398K22.12 (KHDC3L-2)* regulates *KCNQ5* in *cis*

Next, we investigated in detail *RP11-398K22.12* (ENSG00000229852) where the knockdowns by two independent ASOs (ASO_03, ASO_05) successfully reduced the expression of the target lncRNA (67-82% knockdown efficiency, respectively) and further downregulated its neighboring genes, *KCNQ5* and its divergent partner novel lncRNA *CATG00000088862.1* (Fig. 5A). While the two genomic loci occupy Chromosome 6 and are 650kb away, Hi-C analysis (Supplemental Methods; Supplemental Fig. S7) showed that they are located within the same topologically associated domain (TAD) and spatially co-localized (Fig. 5B). Moreover, chromatin-enrichment and single molecule RNA-FISH of *RP11-398K22.12* (Fig. 5C) suggested its highly localized *cis*-regulatory role.

**Figure 5.**
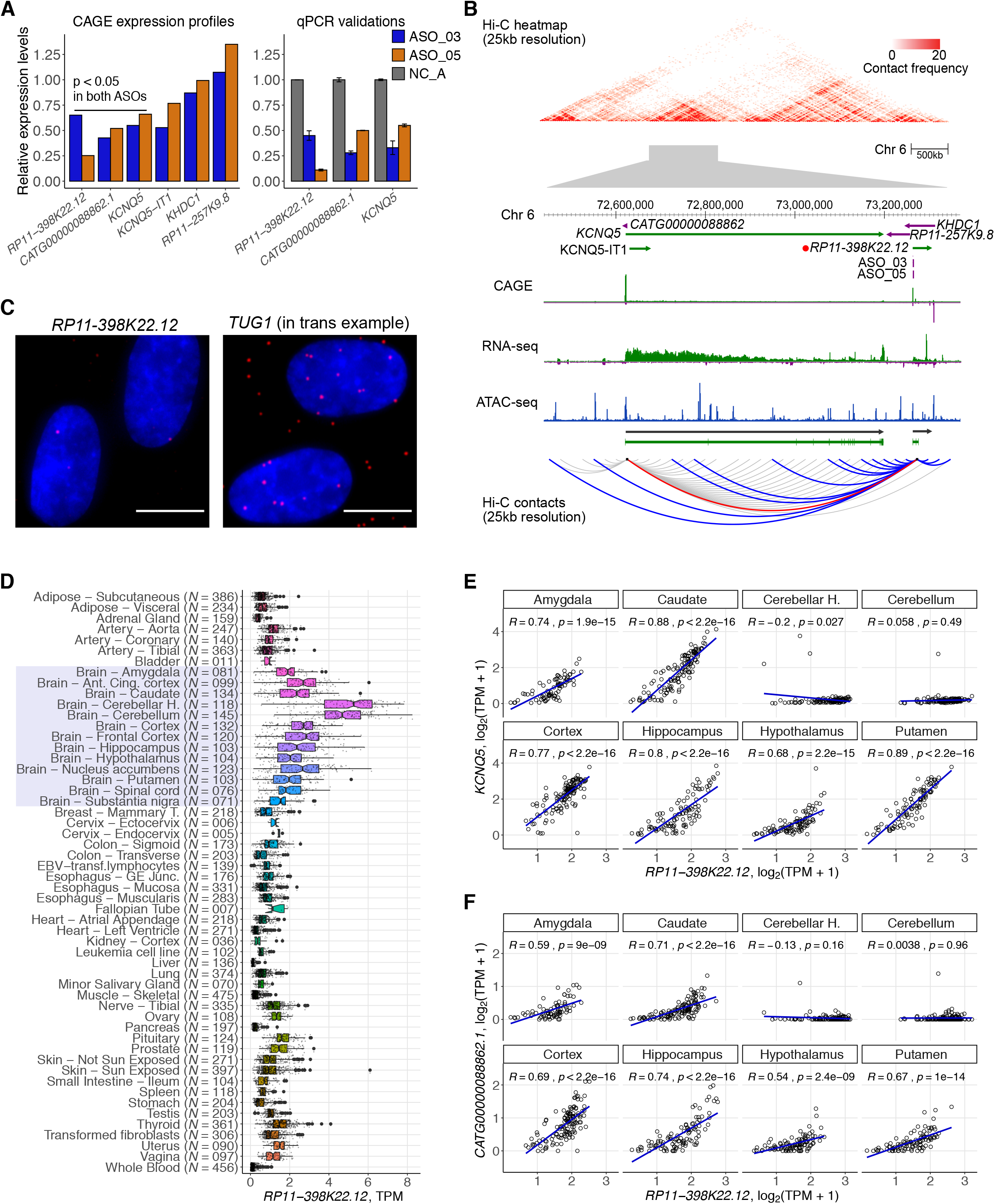
*RP11-398K22.12* down-regulates *KCNQ5* and *CATG00000088862.1* in cis. (*A*) Changes in expression levels of detectable genes in the same topologically associated domain (TAD) as *RP11-398K22.12* based on Hi-C analysis. Both *KCNQ5* and *CATG00000088862.1* are down-regulated (p < 0.05) upon the knockdown of *RP11-398K22.12* by two independent ASOs in CAGE analysis (left) as further confirmed with RT-qPCR (right). (*B*) (top) Representation of the chromatin conformation in the 4Mb region proximal to the TAD containing *RP11-398K22.12,* followed by the locus gene annotation, CAGE, RNA-seq and ATAC-seq data for native HDFs. (bottom) Schematic diagram showing Hi-C predicted contacts of *RP11-398K22.12* (blue) and *KCNQ5* (grey) (25Kb resolution, frequency >= 5) in HDF cells. Red line indicates *RP11-398K22.12* and *KCNQ5* contact. (*C*) FISH image for *RP11-398K22.12* suggesting proximal regulation. *TUG1* FISH image (suggesting trans regulation) is included as a comparison; (bar =10 μm). *(D)* GTEx atlas across 54 tissues *(N* = 9,662 samples) shows relatively high expression levels of *RP11-398K22.12* in 13 distinct brain regions samples (highlighted). *(E)* Expression correlation for *RP11-398K22.12* and *KCNQ5* in 8 out of 13 distinct brain regions, as highlighted in D. *(F)* Expression correlation for *RP11-398K22.12* and *CATG00000088862.1* in 8 out of 13 distinct brain regions, as highlighted in D.

In FANTOM5 (Hon et al. 2017), expression levels of *RP11-398K22.12, KCNQ5* and *CATG00000088862.1* were enriched in brain and nervous system samples, while GTEx (GTEx Consortium 2015) showed their highly-specific expression in the brain, particularly in the cerebellum and the cerebellar hemisphere (Fig. 5D). GTEx data also showed that expression of *RP11-398K22.12* with *KCNQ5* and *CATG00000088862.1* was highly correlated across neuronal tissues (Fig. 5E,F), with the exception of cerebellum and cerebellar hemisphere, potentially due to relatively lower levels of *KCNQ5* and *CATG00000088862.1* while levels of *RP11-398K22.12* remained relatively higher. Additionally, we found an eQTL SNP (rs14526472) overlapping with *RP11-398K22.12* and regulating expression of *KCNQ5* in brain caudate (p = 4.2 × 10^-6^; normalized effect size −0.58). All these findings indicate that *RP11-398K22.12* is implicated in the nervous system by maintaining the expression of *KCNQ5* and *CATG00000088862.1* in a *cis*-acting manner.

## Discussion

This study systematically annotates lncRNAs through molecular and cellular phenotyping by selecting 285 lncRNAs from human dermal fibroblasts across a wide spectrum of expression, conservation levels and subcellular localization enrichments. Using ASO technology allowed observed phenotypes to be associated to the lncRNA transcripts, while in contrast CRISPR-based approaches may synchronically influence the transcription machinery at the site of the divergent promoter or affect regulatory elements of the targeted DNA site. Knockdown efficiencies obtained with ASOs were observed to be independent of lncRNA expression levels, subcellular localization, and of their genomic annotation, allowing us to apply the same knockdown technology to various classes of lncRNAs.

We investigated the *cis*-regulation of nearby divergent promoters, which has been reported as one of the functional roles of lncRNA (Luo et al. 2016). However, in agreement with previous studies (Guttman et al. 2011) we did not observe general patterns in the expression response of divergent promoters (Supplemental Fig. S3B). Recent studies suggest that transcription of lncRNA loci that do not overlap with other transcription units may influence RNA polymerase II occupancy on neighboring promoters and gene bodies (Engreitz, Haines, et al. 2016), (Cho et al. 2018). Thus, it is plausible that transcription of targeted lncRNA was maintained, despite suppression of mature or nascent transcripts using ASOs. This further suggests that the functional responses described in this study are due to interference of processed transcripts present either in the nucleus, the cytoplasm or both. While it is arguable that ASOs may interfere with general transcription by targeting the 5’-end of nascent transcripts and thus releasing RNA polymerase II followed by exonuclease-mediated decay and transcription termination (aka “torpedo model”; (Proudfoot 2016)), most of the ASOs were designed across the entire length of the transcript. Since we did not broadly observe dysregulation in nearby genes, interference of transcription or splicing activity is less likely to occur.

We observed a reduction in cell growth for ~7.7% of our target lncRNA genes, which is in-line with previous experiments using CRISPRi-pooled screening, which reported 5.9% (in iPS cells) of lncRNAs exhibiting a cell growth phenotype (Liu et al. 2017). While these rates are much lower than for protein-coding genes (Sokolova et al. 2017), recurrent observations of cell growth (including cell death) phenotypes strongly suggest that a substantial fraction of lncRNAs play an essential role in cellular physiology and viability. Further, when applying image-based analysis, we found that lncRNAs affect cell morphologies (Fig. 2G), which has not been so far thoroughly explored.

Several lncRNAs such as *MALAT1, NEAT1,* and *FIRRE* have been reported to orchestrate transcription, RNA processing, and gene expression (Kopp and Mendell 2018), but are not essential for mouse development or viability. These observations advocate for assays that can comprehensively profile the molecular changes inside perturbed cells. Therefore, in contrast to cell-based assays, functional elucidation via molecular phenotyping provides comprehensive information that cannot be captured by a single phenotypic assay. Herein, the number of overlapping differentially expressed genes between 2 ASOs of the same lncRNA targets, indicated that 10.9% of lncRNAs exert a reproducible regulatory function in HDF.

Although the features of selected lncRNAs being generally similar to those of other lncRNAs expressed in HDF (Fig. 1B-D), the cell type specific nature of lncRNAs and the relatively small sampling size (119 lncRNAs with knockdown transcriptome profiles) used in our study may not fully represent the whole extent of lncRNA in other cell types. However, lncRNA targets that did not exhibit a molecular phenotype may be biologically relevant in other cell types or cell states (Li and Chang 2014); (Liu et al. 2017). At the same time, our results showed that particular lncRNAs expressed broadly in other tissues (e.g., in the human brain) were functional in HDF (in case of *RP11-398K22.12).* Although the exact molecular mechanisms of *RP11-398K22.*12 are not yet fully understood, its potential role in HDF suggests that lncRNAs may be functionally relevant across multiple tissues in spite of the cell-type-specific expression of lncRNAs.

Further, we used siRNA technology to knockdown lncRNA targets as a method for independent validation. When comparing the transcriptomes perturbed by ASOs and siRNAs, concordance was observed only for 3 out of 9 lncRNAs. This discrepancy is likely due to different modes of actions of the two technologies. While ASOs invoke RNase H-mediated cleavage, primarily active in the nucleus, the siRNAs use RNA-inducing silencing complex (RISC) mainly active in the cytoplasm. LncRNAs are known to function in specific subcellular compartments (Chen 2016) and their maturity, secondary structures, isoforms and functions could be vastly different across compartments (Johnsson et al. 2013). Since the majority functional lncRNA are reported to be inside the nucleus (Palazzo and Lee 2018), (Sun et al. 2018), ASO-mediated knockdowns, which mainly target nuclear RNAs, are generally more suitable for functional screenings of our lncRNA (62% found in the nuclear compartment). Besides, the dynamics of secondary effects mediated by different levels of knockdown from different technologies are likely to be observed as discordance when considering the whole transcriptome, where this kind of discordance has been reported previously (Stojic et al. 2018). In contrast, in the MKI67 assay where only a single feature such as growth phenotype is assayed, siRNA knockdown revealed higher reproducibility with ASO knockdown. This suggested that the growth phenotype might be triggered by different specific pathways in ASO- and siRNA-knockdowns.

Previous studies suggests that lncRNAs regulate gene expression in *trans* epigenetically, via direct or indirect interaction with regulators such as DNMT1 (Di Ruscio et al. 2013) or by directly binding to DNA (triplex; (Mondal et al. 2015) or other RNA binding proteins (Tichon et al. 2016). Analysis of cellular localization by fractionation followed by RNA-seq and *in situ* hybridization can indicate whether a given lncRNA may act *in trans* by quantifying its abundance in the nuclear soluble fraction as compared to cytoplasm. While most lncRNAs in nuclear soluble fraction may affect pathways associated with chromatin modification, additional experiments to globally understand their interaction partners will elucidate the molecular mechanism behind *trans*-acting lncRNAs (Li et al. 2017); (Sridhar et al. 2017).

In summary, our study highlights the functional importance of lncRNAs regardless of their expression, localization and conservation levels. Molecular phenotyping is a powerful and generally more sensitive to knockdown mediated changes platform to reveal the functional relevance of lncRNAs that cannot be observed based on the cellular phenotypes alone. With additional molecular profiling techniques, such as RNA duplex maps in living cells to decode common structural motifs (Lu et al. 2016), and Oxford Nanopore Technology (ONT) to annotate the full-length variant isoforms of lncRNA (Hardwick et al. 2019), structure-to-functional relationship of lncRNAs may be elucidated further in the future.

## Methods

### Gene Models and lncRNA targets selections

The gene models used in this study were primarily based on the FANTOM CAGE-associated transcriptome (CAT) at permissive level as defined previously (Hon et al. 2017). From this merged assembly, there were ~2,000 lncRNAs robustly expressed in HDF (TPM ≥ 1). However, we selected lncRNA knockdown targets in an unbiased manner to broadly cover various types of lncRNAs (TPM ≥ 0.2). Briefly, we first identified a list of the lncRNA genes expressed in HDF, with RNA-seq expression at least 0.5 fragments per kilobase per million and CAGE expression at least 1 tag per millions. Then we manually inspected each lncRNA locus in ZENBU genome browser for 1) its independence from neighboring genes on the same strand (if any), 2) support from RNA-seq (for exons and splicing junctions) and CAGE data (for TSS) of its transcript models and 3) support from histone marks at TSS for transcription initiation (H3K27ac) and along gene body for elongation (H3K36me3), from Roadmap Epigenomics Consortium (Roadmap Epigenomics Consortium et al. 2015). A representative transcript model, which best represents the RNA-seq signal, was manually chosen from each locus for design of antisense oligonucleotides (ASOs). In total, 285 lncRNA loci were chosen for ASO suppression. Additional controls *(NEAT1,* protein coding genes Supplemental Table S1) were added including *MALAT1* as an experimental control. For details please refer to the Supplemental Methods.

### ASO design

ASOs were designed as RNase H-recruiting locked nucleic acid (LNA) phosphorothioate gapmers with a central DNA gap flanked by 2-4 LNA nucleotides at the 5’ and 3’ ends of the ASOs. For details please refer to the Supplemental Methods.

### Automated cell culturing, ASO transfection and cell harvesting

Robotic automation (Hamilton^®^) was established to provide stable environment and accurate procedural timing control for cell culturing and transfection. In brief, trypsin-EDTA detachment, cell number and viability quantification, cell seeding, transfection and cell harvesting were performed with automation. All transfections were divided into 28 runs at weekly basis. ASO transfection was performed with duplication. In each run, there were 16 independent transfections with ASO negative control A (NC_A, Exiqon) and 16 wells transfected with an ASO targeting *MALAT-1* (Exiqon).

The HDF cells were seeded in 12-well plates with 80,000 cells in each well 24 hours prior to the transfection. A final concentration of 20 nM ASO and 2 μl lipofectamine RNAiMAX (Thermo Fisher Scientific) were mixed in 200 μl Opti-MEM (Thermo Fisher Scientific). The mixture was incubated at room temperature for 5 min and added to the cells, which were maintained in 1 ml complete medium. The cells were harvested 48 hours post-transfection by adding 200 μl RLT buffer from the RNeasy 96 Kit (Qiagen) after PBS washing. The harvested lysates were kept at −80°C. RNA was extracted from the lysate for real time quantitative RT-PCR (Supplemental Methods).

### ASO transfection for real-time imaging

The HDF cells were transfected manually in 96-well plate to facilitate high-throughput real time imaging. The cells were seeded 24 hours before transfection at a density of 5,200 cells per well. A final concentration of 20 nM ASO and 2 μl lipofectamine RNAiMAX (Thermo Fisher Scientific) were mixed in 200 μl Opti-MEM (Thermo Fisher Scientific). After incubating at room temperature for 5 min, 18 μl of the transfection mix was added to 90 μl complete medium in each well. The ASOs were divided in 14 runs and transfected in duplicates. Each plate accommodated 6 wells of NC_A control, 2 wells of *MALAT1* ASO control and 2 wells of mock-transfection (lipofectamine alone) control.

Phase-contrast images of transfected cells were captured every 3 hours for 2 days with 3 fields per well by the IncuCyte^®^ live-cell imaging system (Essen Bioscience). The confluence in each field was analyzed by the IncuCyte^®^ software. The mean confluence of each well was taken along the timeline until the mean confluence of the NC_A control in the same plate reached 90%. The growth rate in each well was calculated as the slope of a linear regression. A normalized growth rate of each replicate was calculated as the growth rate divided by the mean growth rate of the 6 NC_A controls from the same plate. Negative growth rate was derived when cells shrink and/or detach. As these rates of cell depletion could not be normalized by the rate of growth, negative values were maintained to indicate severe growth inhibition. Student’s *t*-test was performed between the growth rate of the duplicated samples and the 6 NC_A controls, assuming equal variance.

### Cell morphology quantification

For each transfection, representative phase-contrast image at a single time point was exported from the Incucyte time-series. These raw images were first transformed to probability maps of cells by pixel classification using ilastik (1.3.2) (Berg et al. 2019). The trained model was then applied to all images where the predicted probability maps of cells (grey scale, 16 bits tiff format) were subsequently used for morphology quantification in CellProfiler (3.1.5) (Carpenter et al. 2006). For details please refer to the Supplemental Methods.

### MKI67 staining upon lncRNA knockdown

For the selected four lncRNA targets showing >25% growth inhibition, we used two siRNAs and ASOs with independent sequences. The transfected cells were fixed by adding pre-chilled 70% ethanol and incubated in −20°C. The cells were washed by FACS buffer (2% FBS in PBS, 0.05% NaN3) twice. FITC-conjugated MKI67 (20Raj1, eBioscience) was applied to the cells and subjected to flow cytometric analysis. Knockdown efficiency by siRNA was determined by realtime quantitative RT-PCR using the same 3 primer pairs as for ASO knockdown efficiency. For details please refer to the Supplemental Methods.

### Wound closure assay

The HDF cells were transfected by 20nM ASO as described earlier in 12-well plates. The cells were re-plated at 24 hours post-transfection into a 96-well ImageLock plate (Essen BioScience) at a density of 20,000 cells per well. At 24 hours after seeding, cells form a spatially uniform monolayer with 95-100% cell confluence. The cells were incubated with 5 μg/mL mitomycin-C for 2 hours to inhibit cell division. Then, medium was refreshed and a uniform scratch was created in each well by the WoundMaker™(Essen BioScience). The closure of the wound was monitored by IncuCyte^®^ live-cell imaging system (Essen Bioscience) every 2 hours for 24 hours. The RNA was harvested after the assay for real-time quantitative RT-PCR. For details please refer to the Supplemental Methods.

#### Cap analysis of gene expression (CAGE)

Four micrograms of purified RNA were used to generate libraries according to the nAnT-iCAGE protocol (Murata et al. 2014). For details please refer to the Supplemental Methods.

#### Chromosome conformation capture (Hi-C)

Hi-C libraries were prepared essentially as described previously (Fraser, Ferrai, et al. 2015; Lieberman-Aiden et al. 2009) with minor changes to improve the DNA yield of Hi-C products (Fraser, Williamson, et al. 2015). For details please refer to the Supplemental Methods.

## Data Access

All raw and processed sequencing data generated in this study have been submitted to the DNA Data Bank of Japan (DDBJ; https://www.ddbj.nig.ac.jp/) under accession numbers (DRA008311, DRA008312, DRA008436, DRA008511) or can be accessed through the FANTOM6 project portal https://fantom.gsc.riken.jp/6/datafiles. The analysis results can be downloaded from https://fantom.gsc.riken.jp/6/suppl/Ramilowski_et_al_2020/data/ and interactively explored using our in-house portal https://fantom.gsc.riken.jp/zenbu/reports/#FANTOM6/.

## Acknowledgements

<u>General</u>: We would also like to thank Linda Kostrencic, Hiroto Atsui, Emi Ito, Nobuyuki Takeda, Tsutomu Saito, Teruaki Kitakura, Yumi Hara, Machiko Kashiwagi, Masaaki Furuno at RIKEN Yokohama for assistance in arranging collaboration agreements, ethics applications, computational infrastructure and the FANTOM6 meetings. The authors wish to acknowledge RIKEN GeNAS for generation and sequencing of the CAGE libraries and subsequent data processing.

## Author Information

Competing interests - all authors declare no competing interest

