## Supplementary methods for "Functional Annotation of Human Long Non-Coding RNAs via Molecular Phenotyping"

**Gene Models and lncRNA targets selection**

Gene models used in this study were primarily based on the FANTOM CAGE-associated transcriptome (CAT) at the permissive level as previously defined [(Hon et al. 2017)](http://sciwheel.com/work/citation?ids=3205874&pre=&suf=&sa=0), with additional *de novo* transcript models constructed from human dermal fibroblasts (HDFs) and induced pluripotent stem cells (iPSCs) RNA sequencing data. In brief, CAGE sequencing was performed on the total RNA, and RNA-seq was performed on ribosomal-RNA depleted RNA, from HDFs and iPSCs. CAGE and RNA-seq reads were mapped onto hg19 using TopHat2 (2.0.12) [(Kim et al. 2013)](http://sciwheel.com/work/citation?ids=396582&pre=&suf=&sa=0) with default parameters. RNA-seq reads were *de novo* assembled for each cell line using Cufflinks as previously described [(Hon et al. 2017)](http://sciwheel.com/work/citation?ids=3205874&pre=&suf=&sa=0) and the transcript models with their 5’ends supported by CAGE reads were retained. LncRNA genes were identified from these retained transcript models as previously described [(Hon et al. 2017)](http://sciwheel.com/work/citation?ids=3205874&pre=&suf=&sa=0). The novel lncRNA genes (i.e. loci non-overlapping with FANTOM CAT) were merged with the permissive FANTOM CAT, and the merged assembly were lifted over [(Hinrichs et al. 2006)](http://sciwheel.com/work/citation?ids=652864&pre=&suf=&sa=0) from hg19 to hg38.

From this merged assembly, we selected lncRNA knockdown targets in an unbiased manner to broadly cover various types of lncRNAs. Briefly, we first identified a list of the lncRNA genes expressed in HDF, with RNA-seq expression levels of at least 0.5 fragments per kilobase per million and CAGE expression of at least 1 tag per millions. We then manually inspected each lncRNA locus in ZENBU genome browser for: i) its independence from neighboring genes on the same strand (if any), ii) support from RNA-seq (for exons and splicing junctions) and CAGE data (for TSS) of its transcript models and iii) support from histone marks at TSS for transcription initiation (H3K27ac) and along the gene body for elongation (H3K36me3), from Roadmap Epigenomics Consortium [(Roadmap Epigenomics Consortium et al. 2015)](http://sciwheel.com/work/citation?ids=48808&pre=&suf=&sa=0). A representative transcript model, which best represents the RNA-seq signal, was manually chosen from each locus to design antisense oligonucleotides (ASOs). In total, 285 lncRNA loci were chosen for ASO suppression. Additional controls (NEAT1, protein coding genes; Supplemental Table S1) were added, including MALAT1 as an experimental control.

**ASO design**

ASOs were designed as RNase H-recruiting locked nucleic acid (LNA) phosphorothioate gapmers with a central DNA gap flanked by 2-4 LNA nucleotides at the 5’ and 3’ ends of the ASOs. For each lncRNA target, we used the unspliced transcript sequence from FANTOM CAT as a template for designing a minimum of 5 ASOs per lncRNA.

Briefly, an exhaustive list of all target recognition sequences of 16-19 nt in length within the unspliced target lncRNA sequence was analyzed against the entire transcriptome to determine the prevalence of each *k*-mer, when allowing for 0 or 1 mismatched positions. Furthermore, the accessibility of each *k*-mer in the target sequence was estimated using RNAplfold using the ViennaRNA package [(Lorenz et al. 2011)](http://sciwheel.com/work/citation?ids=52671&pre=&suf=&sa=0) [[doi:10.1186/1748-7188-6-26](http://dx.doi.org/10.1186/1748-7188-6-26)]. Subsequently, ASO designs were generated by applying various LNA/DNA design motifs to the reverse complement of the *k*-mer sequences. The LNA/DNA design patterns consisted of any combination of 2-4 LNA modifications at the 5’ and 3’ends and central DNA window. For each ASO design, we calculated a series of characteristics, including predicted melting temperature for the ASO:RNA duplex, presence of problematic sequence motifs (G-quadruplex and other homopolymers, 5’ terminal GG) as well as propensity to fold/co-fold estimated by using RNAfold and RNAcofold (ViennaRNA package) on the ASO sequence. Finally, ASOs were selected based on properties of both the *k*-mer binding site in the target lncRNA sequence and characteristics of the different ASO designs, such that ASOs had zero perfectly matched off-targets, and no more than 1 imperfectly matched off-target, when allowing for 1 mismatched position. In addition, ASOs were selected to have a predicted melting temperature (T_m_) in the range of 50-56°C [(Pedersen et al. 2014)](http://sciwheel.com/work/citation?ids=6830318&pre=&suf=&sa=0).

A total of 2,055 ASOs targeting 285 lncRNAs were selected for the study (Supplemental Table S1). The ASOs were synthesized by Exiqon (~1,500 ASOs) and GeneDesign (~500 ASOs) Inc. and subsequently classified as exonic or intronic based on their overlap with exons in FANTOM CAT gene models (available at <https://fantom.gsc.riken.jp/zenbu/gLyphs/#config=wE8kSc2aJ5ZtJzixaOOon>).

**Cell culture**

Human dermal fibroblasts (HDFs) used were primary cells derived from the dermis of normal human neonatal foreskin cells (Lonza, catalog number: C2509). The cells were cultured in Dulbecco's Modified Eagle's medium (high glucose with L-glutamine) supplemented with 10% fetal bovine serum at 37°C in a 5% CO_2_ incubator. The passage number of the cells for transfection was maintained at six or seven.

**RNA purification**

The harvested lysates were subjected to purification using RNeasy 96 Kit (Qiagen) and epMotion automated liquid handling systems (Eppendorf) according to the manufacturer’s instructions, except an additional washing step by RPE buffer. The eluted RNA was qualified and quantified by the Dropsense spectrophotometry platform (Trinean). The RNA was stored at -80°C.

**Real-time quantitative RT-PCR**

Real-time quantitative RT-PCR was performed by One Step SYBR PrimeScript™ RT-PCR Kit II (Takara) using the epMotion automated liquid handling system (Eppendorf). For each sample, three primer pairs against the specific lncRNA target were used. The expression level was normalized by GAPDH while the knockdown efficiency was calculated from the fold-change between each sample and NC_A. The knockdown efficiency of *MALAT-1* was monitored along each run and across all the runs where all the samples have >90% knockdown. Sequences of primers are listed in (Supplemental Table [S2](https://fantom6-collaboration.gsc.riken.jp/webdav/home/jordan/fantom6/HDF/Manuscript/tables/excel/Supplementary_Table_S2.xlsx)). In general, the knockdown RNA samples having >50% knockdown efficiency shown consistently by one of the primer pairs were subjected to CAGE. Exceptions included insufficient amount of RNA due to cell death and great knockdown variation between replicates.

**Cell fractionation for RNA-sequencing**

A previously described method [(Conrad and Ørom 2017)](http://sciwheel.com/work/citation?ids=6718119&pre=&suf=&sa=0) was adopted for the isolation of cytoplasmic, nucleoplasmic and chromatin-associated RNA. Approximately 10 million cells were used per fractionation experiment. Briefly, trypsinized cells were washed and lysed using cold lysis buffer containing 0.15% Igepal CA-630, 10 mM Tris pH 7.5, 150 mM NaCl. The lysate was centrifuged in a sucrose cushion, after which the supernatant was taken as the cytoplasmic fraction. The nuclear pellet was washed once in buffer containing 20 mM HEPES pH 7.5, 50% glycerol, 75 mM NaCl, 1 mM DTT, 0.5 mM EDTA and suspended again in the same buffer. An equal volume of nuclear lysis buffer containing 20 mM HEPES pH 7.5, 300 mM NaCl, 1M Urea, 1% Igepal CA-630, 10 mM MgCl_2_, 1 mM DTT, 0.2 mM EDTA was added and incubated on ice for 5 min. After centrifugation, the supernatant was considered as the nucleoplasmic fraction and the pellet as the chromatin fraction. The chromatin pellet was washed once in buffer containing 10 mM HEPES pH 7.5, 10 mM KCl, 10% glycerol, 340 mM sucrose, 4 mM MgCl_2_, 1 mM DTT and suspended in the same buffer. RNA from each fraction was isolated using Trizol LS (Invitrogen) according to manufacturer’s instructions. To ensure RNA purity, DNase I treatment followed by phenol-chloroform extraction was conducted. RNA isolated from each fraction was subjected to RNA-sequencing.

**MKI67 staining upon lncRNA knockdown**

For the selected four lncRNA targets showing >25% growth inhibition, we used two siRNAs and ASOs with independent sequences. The *Silencer*® *Select siRNAs* were obtained from Invitrogen and ASOs were from geneDesign (Supplemental Table[7](https://fantom6-collaboration.gsc.riken.jp/webdav/home/jordan/fantom6/HDF/Manuscript/tables/excel/Supplementary_Table_S3.xlsx)). The HDF cells were transfected with 20 nM siRNA or ASO in 12-well plates by lipofection. The cells were seeded 24 hours before transfection at a density of 60,000 cells per well. At 48 hours post-transfection, cells were washed by PBS and harvested by trypsin-EDTA. Cells from the two wells with the same transfection were collected into one tube. After PBS washing, the cells were fixed by adding pre-chilled 70% ethanol and incubated in -20°C for at least 2 hours. Ethanol was then removed by centrifugation and the cells were washed by FACS buffer (2% FBS in PBS, 0.05% NaN3) twice. FITC-conjugated MKI67 (20Raj1, eBioscience) was applied to cells at a ratio of 8 µl per 150,000 cells. Same concentration of FITC-conjugated mouse IgG1 kappa antibody (eBioscience) was used as isotypic control. After 1 hour of incubating at 4°C, cells were washed by FACS buffer and subjected to flow cytometric analysis. Knockdown efficiency by siRNA was determined by real-time quantitative RT-PCR using the same 3 primer pairs as for ASO knockdown efficiency.

**Wound closure assay**

The HDF cells were transfected by 20 nM ASO as described earlier in 12-well plates. The cells were re-plated at 24 hours post-transfection into a 96-well ImageLock plate (Essen BioScience) at a density of 20,000 cells per well. At 24 hours after seeding, cells form a spatially uniform monolayer with 95-100% cell confluence. The cells were incubated with 5 µg/mL mitomycin-C for 2 hours to inhibit cell division. Then, medium was refreshed and a uniform scratch was created in each well by the WoundMaker™ (Essen BioScience). After changing the medium twice, the cells were maintained in medium with 0.5% FBS. The condition of mitomycin C and serum concentrations was tested with HDF and showed complete growth inhibition without severe morphological change. The closure of the wound was monitored by IncuCyte® live-cell imaging system (Essen Bioscience) every 2 hours for 24 hours. At each time point, the relative wound density was calculated by the cell confluence within the wound area normalized by the cell confluence of the non-wound area in the same image. The relative wound closure rate was calculated as the slope of the linear regression of the relative wound density against time, followed by normalization with that of NC_A. The RNA was harvested after the assay for real-time quantitative RT-PCR.

**Cell morphology quantification**

For each transfection, representative phase-contrast image at a single time point was exported from the IncuCyte® time-series, when the confluence of NC_A-transfected cells in each batch was around 80% (at 30 – 36 hours post-transfection). These raw images were first transformed to probability maps of cells by pixel classification using ilastik (1.3.2) [(Berg et al. 2019)](http://sciwheel.com/work/citation?ids=7553593&pre=&suf=&sa=0). Three-pixel categories including cell, cell boundary, and background were manually labeled in a set of randomly selected images. The trained model was then applied to all images in the batch mode. The predicted probability maps of cells (grey scale, 16 bits tiff format) were subsequently used for morphology quantification in CellProfiler (3.1.5) [(Carpenter et al. 2006)](http://sciwheel.com/work/citation?ids=172434&pre=&suf=&sa=0). In brief, binarized segmented cell images were obtained using the module IdentifyPrimaryObjects (thresholding by Global Otsu method, followed by intensity-based declumping) and the morphology measurements were performed by the module MeasureObjectSizeShape (<http://cellprofiler-manual.s3.amazonaws.com/CellProfiler-3.0.0/modules/measurement.html#measureobjectsizeshape>). All values (medians) were further normalized by the NC_A values from the matching transfection plate, identically to normalizing the growth rate.

**Cap analysis of gene expression (CAGE)**

Four micrograms of purified RNA were used to generate libraries according to the nAnT-iCAGE protocol [(Murata et al. 2014)](http://sciwheel.com/work/citation?ids=463108&pre=&suf=&sa=0). Briefly, random primer with anchor was used for cDNA synthesis, followed by biotinylation and RNaseI digestion. After cap trapping of the 5′ end complete cDNA and linker ligation, second-strand was synthesized for dsDNA.Libraries were combined in 8-plex using different barcodes and were subjected to 50-base single-end sequencing using an Illumina HiSeq 2500 instrument. Tag were de-multiplexed and mapped to human genome assembly hg38 using TopHat2 (2.0.12). The average mapping rate was 68.9% with around 10 million mapped counts obtained on average across all samples.

The samples with mapped counts lower than 500,000 were excluded from further analysis. Several samples were flagged samples as “questionable” if their mapped counts were less than 1,000,000 or their A260/A230 ratio were less than 1.0. Additionally, we manually flagged as “questionable” if any exceptional QC metrics were detected after sequencing, which includes lower amounts of library volumes and possible errors in experiments.

**CAGE promoter and gene expression and batch correction**

Expression for CAGE promoters was estimated by counting the numbers of mapped tags falling using 379,953 promoter regions of gene models as described in ‘*Gene Models and lncRNA targets selections*’. From there, expression of each of 124,245 genes was estimated by summing up the expression values of all promoters assigned to a given gene.

Batch correction was performed on the log-transformed ‘cpm’ values with the prior.count set to 0.25 and normalized for the library sizes using ‘removeBatchEffect’ function from the ‘limma’ R package where the ‘batch’ was attributed to the CAGE sequencing runs and ‘batch2’, ‘design’ and ‘covariates’ parameters not used.

logCPM <- cpm(dge, log=TRUE, prior.count=0.25, normalized.lib.sizes=T)

removeBatchEffect(logCPM, batch=‘CAGE_lsid’)

**Differential promoter and gene expression**

Differential gene expression was carried separately for each ASO knockdown (mainly duplicated CAGE libraries) against matching (the same CAGE sequencing run) negative controls using DESeq2 (2.16) [(Love et al. 2014)](http://sciwheel.com/work/citation?ids=129353&pre=&suf=&sa=0) with default parameters. For a few ASO knockdowns where CAGE replicates were in distinct sequencing runs, a generalized linear model (*glm*) with appropriate design was used. Only genes with a mean count ≥ 1TPM in either knockdown or negative control libraries were tested. Promoter/gene was differentially expressed if (log_2_FC) > 0.5; FDR ≤ 0.05 and abs(Z-score) > 1.645; where Z-score was obtained by scaling relative expression change (log_2_FC) of each tested gene across all ASO knockdown data.

**Motif Activity Response Analysis (MARA)**

MARA (Suzuki et al. 2009) was performed using batch corrected promoter expression for all the knock-down (KD) and control (both NC_A and NC_B) libraries (970 CAGE libraries). All promoters with expression ≥ 1TPM at least in 70% CAGE libraries (24,014 promoters) were used for the analysis. Transcription factor binding sites (TFBS) for hg38 were predicted as described previously [(Alam et al. 2020)](http://sciwheel.com/work/citation?ids=93652&pre=&suf=&sa=0) using MotEvo [(Arnold et al. 2012)](http://sciwheel.com/work/citation?ids=462788&pre=&suf=&sa=0) for the set of 190 position-weight matrix motifs in SwissRegulon (released on 13 July 2015) [(Pachkov et al. 2013)](http://sciwheel.com/work/citation?ids=3419764&pre=&suf=&sa=0) on a multiple alignment of genome assemblies hg38 (human), rheMac3 (macaque), mm10 (mouse), rn6 (rat), bosTau8 (cow), equCab2 (horse), canFam3 (dog), monDom5 (opossum), and galGal4 (chicken). The number of predicted TFBS were counted for each motif in the -300 to +100 base pair from the midpoint of the FANTOM CAT promoters. Next, MARA was performed to decompose CAGE expression profiles of the promoters in terms of their associated motifs, yielding the activity profile of all the motifs with at least 150 TFBS associated with the expressed promoters across the HDF knockdown and controls samples.

**Gene Set Enrichment Analysis**

Gene set enrichment (GO Biological Process, GO Molecular Function, GO Cellular Component, KEGG, Hallmark and Reactome) analysis was carried separately for each pathway and each ASO knockdown using fgsea (1.8.0) R-package with the following parameters: set.seed(42), minSize=15, maxSize=1000, nperm=100000, nproc=1. Each pathway was tested based on log_2_FC preranked genes values from DESeq2 analysis. Pathway was significantly enriched if abs(NES) > 1; FDR ≤ 0.05 and abs(Z-score)| > 1.645.; where Z-score was obtained by scaling GSEA significance of enrichment: -log_10_(p) × sign(NES) of each tested pathway across all knockdowns.

**FANTOM5 coexpression clusters**

Enrichment of FANTOM5 coexpression clusters was calculated using fgsea (1.8.0) R-package with the following parameters: set.seed(42), minSize=15, maxSize=5000, nperm=100000, nproc=1. Coexpression clusters were tested based on log_2_FC pre-ranked genes values from DESeq2 analysis.

**Cellular and molecular phenotype correlations**

Growth and morphology values were correlated for each ASO against 1) each GSEA pathway using enrichment score, 2) motif activity of each MARA motif (≥1TPM) and 3) each FANTOM5 coexpression clusters using enrichment score.

**Identification of expression response of divergent promoters**

We have defined an antisense divergent partner for a given lncRNA to be a gene that lies on the complementary strand within 2.5kb from the target lncRNA and has an antisense divergent promoter to it. For our analysis, we looked at the 84 of our lncRNAs (targeted by a total of 186 ASOs) with a divergent antisense partner and the significance of their expression changes (DESeq2 FDR ≤ 0.05 and abs(log_2_FC) > 0 criteria).

**One-way ANOVA for the global target growth phenotype**

The one-way ANOVA test was carried out on 194 lncRNAs targets, with 2-10 successful ASOs per target (total of 841 ASOs), using normalized growth rate for each ASO and gave the p-value = 5.01 × 10^-6^. The test was then repeated 100,000 times with lncRNA target labels randomly permuted resulting in the empirical p-value = 0.00037 (Supplemental Fig. S2B). The analysis was done in R 3.5 using the ‘anova’ function (stats 3.6.1 package) and the target labels were permuted using the ‘shuffle’ function (mosaic 1.6.0 package).

**Conditional cutoff for lncRNA target growth phenotype**

The relatively high background growth inhibition (21.1%), which we used to represent the maximum possible rate of growth inhibition mediated by side effects, would introduce bias when there are multiple successful knockdown ASO per target. The background chance of concordant growth inhibition is used to calculate the random chance of the number of concordant ASO along different numbers of ASO assessed using a binomial test (Supplemental Fig. S2C). Briefly, we have taken 21.1% as background growth inhibition, the chance of ≥ 2 ASO in the group of lncRNA with 2 ASO showing successful knockdown is 0.211^2^ = 0.04452, which means false positive happened randomly is less than 5%. We set conditional cutoff for different groups of lncRNA having different numbers of successful knockdown ASO to keep the chance of false positives lower than 5%. At a permissive level (≥ 2 ASO), the number of lncRNA targets showing growth phenotype is 68/194.

**Global analysis of molecular phenotype between ASO-pairs from same and different targets**

In order to determine if ASO knockdown resulted in a specific effect stronger than noise (side effect such as off-target effect or cytotoxicity), we globally analysed all of the transcriptomic profiles with lncRNA targets having at least 2 ASOs (305 KD profiles, 119 lncRNAs) using Jaccard index to quantify the degree of concordance. Briefly, differentially expressed genes from each ASO-knockdown were taken with FDR ≤ 0.05, abs(log_2_FC) > 0.5 and abs(Z-score) > 1.645. The 277 maximum possible pairs were made for the same-target group. Using the same ASOs in these 277 pairs the list was shuffled 10,000 times for permutation. The mean of Jaccard index was calculated for the same target group and for each permutation of the different-target group. The empirical p-value represents the chance that the mean of Jaccard index of the same target group is not greater than the different-target group.

**Identification of concordant ASO-pairs from the same lncRNA targets**

All the ASO-pairs from the same lncRNA targets (maximum possible pairs = 277) were tested for significance to be above the background distribution of Jaccard index. Briefly, the jaccard index of all the ASO-pairs were transformed to Z-score using the mean and standard deviation of the background distribution (different-target ASO-pairs). Using the criteria of Z-score p-value < 0.05 and at least five common DEGs (FDR ≤ 0.05, abs(log_2_FC) > 0.5, abs(Z-score) > 1.645) between the ASO-pair, concordant ASO-pairs were identified as shown in Supplemental Fig. S5B. The overlapping significant DEGs, pathways and MARA motifs of each ASO-pair were summarized and shown in Supplemental Table S5 and Supplemental Fig. S5C for the 16 significant pairs.

**Analysis on concordance between ASO and siRNA knockdown**

Eighteen ASO and eighteen siRNA targeting 9 lncRNA were transfected to HDF using the same protocol described in this study. The RNA samples including 36 knockdowns with duplication, 6 ASO scramble control and 6 siRNA scramble controls were subjected for CAGE sequencing. In DESeq2 we compared each knockdown to their corresponding negative controls (2 VS 6). We did not apply Z-score on the log_2_FC because of the small sampling size. Instead, we required a more stringent cutoff where DEG with FDR ≤ 0.05 and abs(log_2_FC) > 1 were taken. Next, the oligos were paired-up in three different conditions: ASO-ASO, siRNA-siRNA and ASO-siRNA. The oligos were paired for targeting the same lncRNA (ASO-ASO, 9 pairs; siRNA-siRNA, 9 pairs; ASO-siRNA, 36 pairs). Oligos are randomized to target different lncRNA for the same number of pairs with 10,000 permutations. Jaccard index was used to quantify the similarity between oligos from each pair. The empirical p-values represent the probability that the mean Jaccard index of the same-target group is less than that of the different-target group. Concordance was only observed significantly among the ASO modality.

Since we did not observe a global concordance of same-target greater than different-target due to small sampling size, we applied a more stringent cutoff in identifying concordant ASO-siRNA pairs. To do this, we transformed the Jaccard index of each pair into Z-score using the mean and standard deviations from the background (all the different-target pairs). Concordant pairs were identified under the condition where overlapped DEGs ≥ 5 and the Jaccard index significantly greater than the different-target background (p < 0.05).

**Chromosome conformation capture (Hi-C)**

Hi-C libraries were prepared essentially as described previously [(Fraser, Ferrai, et al. 2015; Lieberman-Aiden et al. 2009)](http://sciwheel.com/work/citation?ids=1145036,48455&pre=&pre=&suf=&suf=&sa=0,0) with minor changes to improve the DNA yield of Hi-C products [(Fraser, Williamson, et al. 2015)](http://sciwheel.com/work/citation?ids=761036&pre=&suf=&sa=0). The procedure followed is outlined as a flowchart in Supplemental Fig. S7A, and each step of the protocol is briefly described below.

*Cell cross-linking*

Three biological replicate samples of HDFs (~1×10^7^ cells) grown as described above were fixed at room temperature in media containing 1% formaldehyde (Sigma-Aldrich, catalog number: F8775) for 10 minutes with gentle rocking every 2 minutes. Cross-linking was stopped by quenching the formaldehyde with glycine (Sigma-Aldrich, catalog number: G8898) at a final concentration of 125 mM for 5 minutes at room temperature, followed by 15 minutes on ice. The cells were then scraped off the plates and pelleted by gentle centrifugation at 400 *g* for 10 minutes. The quenched media was removed, and the cell pellets were quick-frozen on dry ice before storage at -80°C.

*Chromatin digestion*

Cell pellets were resuspended into 440 µl of ‘Cold Lysis Buffer’ containing protease inhibitors (10 mM Tris pH 8.0, 10 mM NaCl, 0.2% Igepal; protease inhibitor cocktail as instructed by the manufacturer (Sigma-Aldrich, catalog number: P8340). The cells were incubated on ice for 15 minutes and lysed by twice 20 strokes with a Dounce homogenizer (Pestle B). Cell lysates were centrifuged at 5,000 rpm for 5 minutes at room temperature in a microcentrifuge. The pellets were washed twice with 400 µl of 1X NEBuffer^TM^ 3.1, centrifuged 5 minutes at room temperature at 5,000 rpm, and resuspended in 200 µl of 1X NEBuffer^TM^ 3.1. Each sample was divided equally into 4 tubes before adding 312 µl of 1X NEBuffer^TM^ 3.1 to each tube. 1% SDS (38 µl) was added to each sample and incubated 10 minutes at 65°C. 10% Triton X-100 (44 µl) was mixed into each tube before adding 40 µl of NcoI (10 U/µl; 400 Units; NEB, catalog number: R0193L) and incubating a 37°C overnight.

*Biotin labeling and blunt end ligation*

Digested fragment ends were labeled with biotin by filling-in NcoI 5’ overhangs with Klenow using biotinylated dCTP. Deoxynucleotides (1.5 µl each of 10 mM dATP, dGTP, dTTP, and 37.5 µl of 0.4 mM biotin-14-dCTP (Life Technologies, catalog number: 19518018)) were added to each tube, along with 10 µl of Klenow (5 U/µl; 50 Units; NEB, catalog number: M0210S), and incubated at 37°C for 45 minutes. 10% SDS (86 µl) was mixed into each sample before incubating 30 minutes at 65°C. The samples were transferred to 15 mL conical tubes containing 5.96 mL of water and 1.66 mL of ‘Ligation mix’ (10% Triton X-100 (750 µl), 10X Ligation Buffer (750 µl; 0.5 M Tris pH 7.5, 0.1 M MgCl_2_, 0.1 M DTT), 10 mg/mL BSA (80 µl), 100 mM ATP (80 µl)). T4 DNA ligase (50 µl) was added to each tube (1 U/µl; 50 Units, Invitrogen, catalog number: 15224-025), and incubated 4 hours at 16°C.

*DNA purification*

The DNA was purified by proteinase K digestion followed with phenol/chloroform extraction and ethanol precipitation. Proteinase K (50 µl of 10 mg/mL) was added to each tube and incubated overnight at 65°C. Proteinase K was added again on the next day (50 µl of 10 mg/mL) and incubated for an additional 2 hours. The samples were transferred to 50 mL conical tubes and extracted with 10 mL of phenol by vortexing 2 minutes and centrifugation at 2,465 *g* for 15 minutes at room temperature. The samples were extracted again as above but with phenol/chloroform before precipitating with ethanol. To this end, samples were transferred to 35 mL centrifuge bottles, and their volumes adjusted to 10 mL with 1X TE before 1 mL of 3 M sodium acetate (pH 5.2) was mixed in and 25 mL of ice-cold ethanol was added. Each sample was gently mixed by inversion and incubated 1 hour at -80°C. The DNA was pelleted by centrifugation at 23,281 *g* (13,000 rpm if using SS34 rotor) for 25 minutes at 4°C, washed by resuspending in 10 mL of ice-cold 70% ethanol, and centrifuged again as above. Pellets were each dissolved into 450 µl of 1X TE pH 8.0 and extracted twice with 500 µl of phenol-chloroform by vortexing 1 minute and centrifuging at maximum speed for 5 minutes at room temperature in a microcentrifuge. The samples were precipitated with 3 M sodium acetate pH 5.2 (40 µl) and absolute ethanol (1 mL), incubated at -80°C overnight, and centrifuged at maximum speed for 25 minutes at 4°C. The DNA was washed five times by resuspending in ice-cold 70% ethanol (1 mL), each time centrifuging at maximum speed for 20 minutes at 4°C. The resulting pellets were each dissolved in 25 µl of 1X TE pH 8.0 before the 4 samples from original cell pellets were pooled. 10 mg/mL RNAse A (1 µl; Fermentas, catalog number: EN0531) was added and incubated for 30 minutes at 37°C. The Hi-C DNA was resolved on 0.8% agarose gel containing ethidium bromide (0.5 µg/mL) for qualitative assessment of digestion, ligation, and yield based on calibrated molecular weight markers (Supplemental Fig. S7B).

*Quality control*

The quality of Hi-C DNA was quantitatively assessed by PCR titration of an expected Hi-C contact and digestion of the PCR product as previously outlined [(Lieberman-Aiden et al. 2009)](http://sciwheel.com/work/citation?ids=48455&pre=&suf=&sa=0). The Hi-C DNA was serially diluted in water from 1 to ~0.0005 µL (2-fold; 11 dilutions), and 4 µL aliquots were PCR-amplified (35 cycles) with primers against a gene desert sequence (GD09 NcoI 5’-GCAATTAGTGCTATGCCCATGTTTCCTTTGTTCC-3’, GD10 NcoI 5’-CAGTCTTCTACCGCTCTTGTAATGGGGTT-3’). Half of each PCR reaction was resolved on 1.5% agarose gels containing ethidium bromide (0.5 µg/mL) to verify the presence of amplification products. The remaining products of the first 5 PCR reactions from each titration were pooled, and divided equally into 4 tubes to digest with either NcoI, NsiI, NcoI/NsiI, or buffer control as described previously [(Lieberman-Aiden et al. 2009)](http://sciwheel.com/work/citation?ids=48455&pre=&suf=&sa=0). Digestion reactions were incubated 1 hour at 37°C, and resolved on 2% agarose gels containing ethidium bromide (0.5 µg/mL) to verify greater digestion efficiency with NsiI. In all 3 HDF biological replicates, over 75% of this PCR product digested specifically with NsiI (Supplemental Fig. S7C) pointing to efficient Hi-C product formation in the samples.

*Removal of biotin from unligated ends and DNA shearing*

Biotin at unligated restriction fragment ends was removed with T4 DNA polymerase as we described previously [(Fraser, Ferrai, et al. 2015)](http://sciwheel.com/work/citation?ids=1145036&pre=&suf=&sa=0). Briefly, 10 samples each of 5 µg DNA from individual HDF replicates were mixed with 10µl of 10X NEBuffer^TM^ 2.1, 1 µl of 10 mM dCTP, and 5 Units of T4 DNA polymerase (NEB, catalog number: M0203L) in a final volume of 100 µl in MAXYMum Recovery^TM^ PCR tubes (200 µl; Axygen, part number: PCR-02-L-C). The reactions were incubated in a thermocycler for 2 hours at 12°C, after which the enzyme was inactivated at 75°C for 20 minutes. The samples were transferred to 1.7 mL tubes (Corning™ Costar™ Low Binding Plastic Microcentrifuge Tubes, part # C3207), and the DNA was precipitated by mixing 10 µl of 3 M sodium acetate (pH 5.2) in each sample, followed by 275 µl of ice-cold ethanol, and incubating at -80°C overnight. The DNA was pelleted by centrifugation at 15,000 rpm in a microcentrifuge at 4°C for 25 minutes, washed twice with 500 µl ice-cold 70% ethanol each time centrifuging as above, and the resulting pellets were each dissolved in 130 µl of 10 mM Tris pH 8.5. The DNA was sheared to ~350 bp fragments by sonication with a ‘Covaris M220 Focused-utrasonicator’ using preset settings (DNA_0300_bp_130µl_Snap_Cap_Micro_TUBE). Samples were individually transferred to 1.7 mL tubes and shearing to appropriate size range was verified by resolving 400 ng of each sample onto 1.5% agarose gel containing ethidium bromide (0.5 µg/mL; Supplemental Fig. S7D.

*DNA size selection*

The sheared DNA was size-selected using AMPure XP Beads (Agencourt XP) with ratios of 0.6X and 0.85X according to the manufacturer’s instructions (Beckman Coulter, catalog number: A63880). Briefly, 78 µl of beads was added to each 130 µl sample and incubated with rotation for 30 minutes at room temperature. The samples were placed on a DynaMag^TM^ Magnet (Thermo Fisher Scientific) for 2 minutes, and the supernatant was re-extracted with 32.5 µl of beads as above to capture the appropriate size fragments. The beads were washed twice with 500 µl of ice-cold ethanol and air-dried 5 minutes. The DNA was eluted with 60 µl of 10 mM Tris pH 8.5 by resuspending ten times with a pipette, and the 10 supernatants from each biological replicate were pooled to a 1.7 mL tube after placing samples on the magnetic stand for 2 minutes. The sample volumes were then reduced to 300 µl with a SpeedVac concentrator, and 5 µl of each sample was used to measure DNA concentrations with the Quant-iT^TM^ PicoGreen^TM^ dsDNA Assay Kit as per the manufacturer’s instructions (Thermo Fisher Scientific, catalog number: P11496). This measure will be used below to calculate the amount of sequencing adaptor required. DNA recovery was verified by resolving 5 µl of each sample on a 1.5% agarose gel containing ethidium bromide (0.5 µg/mL; Supplemental Fig. S7E).

*Streptavidin pull-down and DNA end repair*

The biotinylated Hi-C products were pull-down on Dynabeads MyOne^TM^ Streptavidin C1 Beads (Thermo Fisher Scientific, catalog number: 65001) to enhance DNA end repair, 3’ end adenylation, and ligation of sequencing adaptors. The beads (60 µl for each Hi-C biological replicate) were washed with 400 µl ‘Tween Wash Buffer’ (TWB; 5 mM Tris-HCl pH 8.0, 0.5 mM EDTA, 1 M NaCl, 0.05% Tween 20) twice for 3 minutes at room temperature with rotation, and pelleted on a DynaMag^TM^ Magnet for 2 minutes between washes. Beads were next washed once with 300 µl of ‘2X Binding Buffer’ (2XBB; 10 mM Tris-HCl pH 8.0, 1 mM EDTA, 2 M NaCl), and resuspended in 300 µl of 2XBB. Each 300 µl-Hi-C library was added to a tube containing beads and incubated at room temperature for 15 minutes with rotation to bind the DNA. The supernatant was removed, and the beads were washed twice with 400µl of 1XBB (5 mM Tris-HCl pH 8.0, 0.5 mM EDTA, 1 M NaCl), and once with 100 µl of 1X Ligation Buffer (diluted from 10X, NEB, catalog number: B0202S). The supernatant was removed and 100 µl of ‘DNA End Repair Mix’ was added to each sample (10X Ligation Buffer (10 µl), 10 mM dNTPs (4 µl), T4 DNA Polymerase (5µl of 3 U/µl, NEB), T4 Polynucleotide Kinase (5 µl of 10 U/µl, NEB), Klenow (1 µl of 5 U/µl, NEB), water (75 µl). The samples were transferred to MAXYMum Recovery^TM^ 200 µl PCR tubes and placed in a thermocycler for 30 minutes at 20°C. Samples were then transferred to 1.7 mL tubes, and the beads were washed twice with 200 µl of TWB, and twice with 200 µl Elution Buffer (EB; 10 mM Tris-HCl pH 8.5), each time pelleting beads on a magnetic stand for 2 minutes.

*3’ end adenylation and sequencing adapter ligation*

To prevent concatenation of Hi-C products, DNA fragments were 3’ adenylated and ligated to Illumina Paired-End (PE) sequencing adaptors with 3’-T overhangs. To this end, the supernatant was first removed from the beads and 50 µl of 3’ Adenylation Mix’ was added to each tube (10X NEBuffer^TM^ 2 (5 µl), 10 mM dATP (1 µl), Klenow Fragment (3’→5' exo-; 3 µl of 5 U/µl, NEB, catalog number: M0212S), water (41 µl). The samples were transferred to MAXYMum Recovery^TM^ 200 µl PCR tubes and incubated 60 minutes at 37°C in a thermocycler. The samples were transferred to 1.7 mL tubes, and the beads were washed twice with 200 µl of TWB, and twice with 200 µl EB, each time pelleting beads on a magnetic stand for 2 minutes.

The amount of Illumina PE Adaptor needed for ligation depends on the amount of DNA in Hi-C samples. As a rule, 6 pmol of Illumina PE Adaptor (TruSeq^TM^ DNA PCR-Free LT Library Prep Kit-Set A, Illumina, catalog number: FC-121-3001) was used for every 1 µg of DNA measured with the Quant-iT^TM^ PicoGreen^TM^ dsDNA Assay Kit after size selection. The EB was removed from the beads and 45 µl of ‘Adaptor Ligation Mix’ (5X Invitrogen Ligation buffer (10 µl), Illumina PE Adaptors, water) was added. Sample were mixed by pipetting, and 5 µl of T4 DNA Ligase (1 Weiss U/µl, Invitrogen, catalog number: 15224-025) was added to each tube. The samples were transferred to MAXYMum Recovery^TM^ 200 µl PCR tubes and incubated 60 minutes at 20°C in a thermocycler. The samples were transferred to 1.7 mL tubes, and the beads were washed twice for 6 minutes with 400 µl of TWB, and twice with 400 µl EB, each time pelleting beads on a magnetic stand for 1 minute. Each sample was finally resuspended in 50 µl of EB.

*PCR amplification of Hi-C libraries*

As a rule, the lowest possible PCR cycle number should be used to amplify enough Hi-C libraries to reduce amplification biases. To identify the optimal number of PCR cycles, 3 PCR reactions (25 µl) were prepared for each Hi-C library and amplified through either 6, 8, or 10 cycles. Individual PCR reactions were composed of 1 µl Hi-C library, 1.25 µl each of 10 µM Illumina PE 1.0 and PE 2.0 primers, 12.5 µl of Phusion High-Fidelity 2X Master Mix (NEB, catalog number: M0531S), and 9 µl of water. The reactions were conducted in MAXYMum Recovery^TM^ 200 µl PCR tubes using the following PCR program: 1 cycle of 30 seconds at 98°C, either 6, 8, or 10 cycles of 10 seconds at 98°C / 30 seconds at 65°C / 30 seconds at 72°C, and a final cycle of 7 minutes at 72°C. The reactions were resolved on a 2.0% agarose gel containing ethidium bromide (0.5 µg/mL) to select the lowest possible cycle number when a product is detected (Supplemental Fig. S7F).

Eight PCR cycles was used for large-scale amplification of the libraries for which one set of 10 PCR reactions for each replicate were prepared as described above except that 1.5 µl of Hi-C library was used in each reaction. The 10 PCRs from each replicate were pooled and their combined volume was adjusted to 215 µl with water. The amplified DNA was then purified using AMPure XP Beads with a ratio of 0.8X by adding 172 µl of washed beads to each pooled reaction tube and incubating for 10 minutes at room temperature while mixing. The captured DNA was pelleted on a magnetic stand 2 minutes and washed twice on the beads with 1 mL of ice-cold 70% ethanol. The beads were air-dried for no more than 5 minutes and the DNA was eluted from the beads by adding 33 µl of 10 mM Tris pH 8.0, 0.1 mM EDTA and pipetting to resuspend the beads. Sample concentrations were measured using 2 µl of the library and the Quant-iT^TM^ PicoGreen^TM^ dsDNA Assay Kit as per the manufacturer’s instructions (Thermo Fisher Scientific). DNA recovery and quality was verified by resolving 5 µL of each sample on a 2.5% agarose gel containing ethidium bromide (0.5 µg/mL; Supplemental Fig. S7G).

*Sequencing and processing*

The libraries were sequenced on a HiSeq 2500 Illumina platform, and paired-end reads were mapped to hg38 and processed using HiCUP pipeline ver. 0.5.10 [(Wingett et al. 2015)](http://sciwheel.com/work/citation?ids=1106951&pre=&suf=&sa=0). Ditags were mapped against the human genome assembly hg38. Experimental artifacts such as circularized, re-ligated, continuous and incorrect size fragments were filtered out. PCR duplicates were removed from the aligned data. The sequencing and mapping metrics are in (Supplemental Table [S8](https://fantom6-collaboration.gsc.riken.jp/webdav/home/jordan/fantom6/HDF/Manuscript/tables/excel/Supplementary_Table_S3.xlsx)). Processed mapped ditags from all the three replicates were merged using Samtools ver. 1.3.1 [(Li et al. 2009)](http://sciwheel.com/work/citation?ids=48787&pre=&suf=&sa=0) for the further analysis. Significant co-localized regions at 10kb resolution were identified using BioConductor package GOTHiC [(Mifsud et al. 2017)](http://sciwheel.com/work/citation?ids=6684080&pre=&suf=&sa=0). All the identified co-localized regions with p-value ≤ 0.01 and FDR ≤ 0.05 were used for the downstream analysis. TADs were identified using Arrowhead from Juicer pipeline [(Durand et al. 2016)](http://sciwheel.com/work/citation?ids=4967221&pre=&suf=&sa=0). For the downstream analysis FANTOM CAT promoters were mapped to the colocalized regions and TADs.

**RNA Fluorescence in situ hybridization (FISH)**

Oligonucleotide probes against target RNA were designed using the Stellaris Probe Designer version 4.2 (Biosearch Tech). Probes were flanked on both ends with overhang arms serving as annealing sites for secondary probes labeled with a fluorescent dye (Chen et al 2015). Overhang sequences were identical on both ends. Secondary probe sequences have been previously described (Moffit et al 2016) and were labeled on the 3’ end with Atto 647N. All probe sequences used in this study can be found in (Supplemental Table S9). Two-step hybridization was performed using a previously-described procedure [(Kouno et al. 2019)](http://sciwheel.com/work/citation?ids=6718121&pre=&suf=&sa=0). Briefly, fibroblasts were seeded onto coverslips overnight and fixed in 4% formaldehyde in PBS for 10 min at room temperature. Following fixation, coverslips were treated twice with ice-cold 0.1% sodium borohydride for 5 min at 4°C. Coverslips were washed three times in PBS, followed by cell permeabilization in 0.5% Triton X-100 in PBS for 10 min at room temperature. Coverslips were again washed three times in PBS and treated with 70% formamide in 2X SSC for 10 min at room temperature. Coverslips were washed twice in ice-cold PBS and once in ice-cold 2X SSC. Coverslips were either used immediately for hybridization or stored in 70% ethanol for no longer than a week, in which case were washed in PBS twice and once in 2X SSC at room temperature prior to hybridization. For hybridization, coverslips were incubated at 37°C overnight in hybridization buffer containing 252 mM primary probe inside a humid chamber. Hybridization buffer contained 10% dextran sulfate, 1 ug/ul yeast tRNA, 2 mM vanadyl ribonucleoside complex, 0.02% BSA, 10% formamide, 2X SSC. Excess probe was removed by two washes for 30 min at room temperature in wash buffer containing 30% formamide, 2X SSC, 0.1% Triton X-100, followed by a rinse in 2X SSC. For second hybridization, coverslips were incubated at 37°C for 1.5 hours in hybridization buffer containing 30 nM secondary probe. Excess probe was washed twice for 20 min at room temperature. Coverslips were stained for 5 min in 2 mg/mL Hoescht in PBS, washed three times in PBS, and mounted on a glass slide with SlowFade Gold Antifade Mountant (Invitrogen). Imaging was done on a DeltaVision microscope (GE Healthcare) equipped with a sCMOS sensor. Image processing was done using FIJI [(Chen et al. 2015; Moffitt et al. 2016; Kouno et al. 2019)](http://sciwheel.com/work/citation?ids=93458,3435052,6718121&pre=&pre=&pre=&suf=&suf=&suf=&sa=0,0,0)

[**References**](http://sciwheel.com/work/bibliography)

Alam, T., Agrawal, S., Severin, J., Young, R., et al. 2020. Comparative transcriptomics of primary cells in vertebrates, *Genome Research* (in press)

[Arnold, P., Erb, I., Pachkov, M., Molina, N. and van Nimwegen, E. 2012. MotEvo: integrated Bayesian probabilistic methods for inferring regulatory sites and motifs on multiple alignments of DNA sequences. *Bioinformatics* 28(4), pp. 487–494.](http://sciwheel.com/work/bibliography/462788)

[Berg, S., Kutra, D., Kroeger, T., et al. 2019. ilastik: interactive machine learning for (bio)image analysis. *Nature Methods* 16(12), pp. 1226–1232.](http://sciwheel.com/work/bibliography/7553593)

[Carpenter, A.E., Jones, T.R., Lamprecht, M.R., et al. 2006. CellProfiler: image analysis software for identifying and quantifying cell phenotypes. *Genome Biology* 7(10), p. R100.](http://sciwheel.com/work/bibliography/172434)

[Chen, K.H., Boettiger, A.N., Moffitt, J.R., Wang, S. and Zhuang, X. 2015. RNA imaging. Spatially resolved, highly multiplexed RNA profiling in single cells. *Science* 348(6233), p. aaa6090.](http://sciwheel.com/work/bibliography/93458)

[Conrad, T. and Ørom, U.A. 2017. Cellular Fractionation and Isolation of Chromatin-Associated RNA. *Methods in Molecular Biology* 1468, pp. 1–9.](http://sciwheel.com/work/bibliography/6718119)

[Durand, N.C., Shamim, M.S., Machol, I., et al. 2016. Juicer Provides a One-Click System for Analyzing Loop-Resolution Hi-C Experiments. *Cell Systems* 3(1), pp. 95–98.](http://sciwheel.com/work/bibliography/4967221)

[Fraser, J., Ferrai, C., Chiariello, A.M., et al. 2015. Hierarchical folding and reorganization of chromosomes are linked to transcriptional changes in cellular differentiation. *Molecular Systems Biology* 11(12), p. 852.](http://sciwheel.com/work/bibliography/1145036)

[Fraser, J., Williamson, I., Bickmore, W.A. and Dostie, J. 2015. An Overview of Genome Organization and How We Got There: from FISH to Hi-C. *Microbiology and Molecular Biology Reviews* 79(3), pp. 347–372.](http://sciwheel.com/work/bibliography/761036)

[Hinrichs, A.S., Karolchik, D., Baertsch, R., et al. 2006. The UCSC Genome Browser Database: update 2006. *Nucleic Acids Research* 34(Database issue), pp. D590-8.](http://sciwheel.com/work/bibliography/652864)

[Hon, C.-C., Ramilowski, J.A., Harshbarger, J., et al. 2017. An atlas of human long non-coding RNAs with accurate 5’ ends. *Nature* 543(7644), pp. 199–204.](http://sciwheel.com/work/bibliography/3205874)

[Kim, D., Pertea, G., Trapnell, C., Pimentel, H., Kelley, R. and Salzberg, S.L. 2013. TopHat2: accurate alignment of transcriptomes in the presence of insertions, deletions and gene fusions. *Genome Biology* 14(4), p. R36.](http://sciwheel.com/work/bibliography/396582)

[Kouno, T., Moody, J., Kwon, A.T.-J., et al. 2019. C1 CAGE detects transcription start sites and enhancer activity at single-cell resolution. *Nature Communications* 10(1), p. 360.](http://sciwheel.com/work/bibliography/6718121)

[Lieberman-Aiden, E., van Berkum, N.L., Williams, L., et al. 2009. Comprehensive mapping of long-range interactions reveals folding principles of the human genome. *Science* 326(5950), pp. 289–293.](http://sciwheel.com/work/bibliography/48455)

[Li, H., Handsaker, B., Wysoker, A., et al. 2009. The Sequence Alignment/Map format and SAMtools. *Bioinformatics* 25(16), pp. 2078–2079.](http://sciwheel.com/work/bibliography/48787)

[Lorenz, R., Bernhart, S.H., Höner Zu Siederdissen, C., et al. 2011. ViennaRNA Package 2.0. *Algorithms for Molecular Biology* 6, p. 26.](http://sciwheel.com/work/bibliography/52671)

[Love, M.I., Huber, W. and Anders, S. 2014. Moderated estimation of fold change and dispersion for RNA-seq data with DESeq2. *Genome Biology* 15(12), p. 550.](http://sciwheel.com/work/bibliography/129353)

[Mifsud, B., Martincorena, I., Darbo, E., et al. 2017. GOTHiC, a probabilistic model to resolve complex biases and to identify real interactions in Hi-C data. *Plos One* 12(4), p. e0174744.](http://sciwheel.com/work/bibliography/6684080)

[Moffitt, J.R., Hao, J., Wang, G., Chen, K.H., Babcock, H.P. and Zhuang, X. 2016. High-throughput single-cell gene-expression profiling with multiplexed error-robust fluorescence in situ hybridization. *Proceedings of the National Academy of Sciences of the United States of America* 113(39), pp. 11046–11051.](http://sciwheel.com/work/bibliography/3435052)

[Murata, M., Nishiyori-Sueki, H., Kojima-Ishiyama, M., Carninci, P., Hayashizaki, Y. and Itoh, M. 2014. Detecting expressed genes using CAGE. *Methods in Molecular Biology* 1164, pp. 67–85.](http://sciwheel.com/work/bibliography/463108)

[Pachkov, M., Balwierz, P.J., Arnold, P., Ozonov, E. and van Nimwegen, E. 2013. SwissRegulon, a database of genome-wide annotations of regulatory sites: recent updates. *Nucleic Acids Research* 41(Database issue), pp. D214-20.](http://sciwheel.com/work/bibliography/3419764)

[Pedersen, L., Hagedorn, P.H., Lindholm, M.W. and Lindow, M. 2014. A Kinetic Model Explains Why Shorter and Less Affine Enzyme-recruiting Oligonucleotides Can Be More Potent. *Molecular therapy. Nucleic acids* 3, p. e149.](http://sciwheel.com/work/bibliography/6830318)

[Roadmap Epigenomics Consortium, Kundaje, A., Meuleman, W., et al. 2015. Integrative analysis of 111 reference human epigenomes. *Nature* 518(7539), pp. 317–330.](http://sciwheel.com/work/bibliography/48808)

Suzuki, H., Forrest, A.R.R., van Nimwegen, E., et al. 2009. The Transcriptional Network That Controls Growth Arrest and Differentiation in a Human Myeloid Leukemia Cell Line. *Nature Genetics* 41(5), pp. 553–562.

[Wingett, S., Ewels, P., Furlan-Magaril, M., et al. 2015. HiCUP: pipeline for mapping and processing Hi-C data. [version 1; peer review: 2 approved, 1 approved with reservations]. *F1000Research* 4, p. 1310.](http://sciwheel.com/work/bibliography/1106951)
